# Characterising cancer-stroma interactions through high-content phenotyping from microscopy time-lapses

**DOI:** 10.1101/2025.10.02.680037

**Authors:** Laura Wiggins, Jodie R. Malcolm, Karen Hogg, Peter J. O’Toole, Julie Wilson, William J. Brackenbury

## Abstract

Understanding how cancer-stromal interactions shape cancer progression requires tools that can capture dynamic phenotypic changes in physiologically relevant conditions. Traditional approaches for studying co-culture interactions, such as transcriptomics and flow cytometry, provide valuable insights but are limited by their static nature and reliance on fixed or dissociated cells. In contrast, label-free time-lapse microscopy preserves temporal and spatial context, enabling observation of live-cell behaviours over time. A major challenge, however, lies in the analysis of the resulting high-dimensional datasets. Using co-cultures of breast cancer cells and cancer-associated fibroblasts (CAFs) as a model system, we show that the CellPhe toolkit enables label-free identification and phenotypic characterisation of different cell types within complex live-cell imaging datasets. Our analysis shows that exposure to CAFs drives marked phenotypic shifts in breast cancer cells, including elongation, loss of cell–cell adhesion, and redistribution of intracellular components - hallmarks of epithelial–mesenchymal transition (EMT). To probe the underlying mechanisms, we performed a Luminex immunoassay on CAF-conditioned media and identified secreted analytes strongly associated with EMT induction. Together, these results highlight how automated phenotyping can be integrated with molecular profiling to identify and characterise cellular processes shaped by stromal interactions and reveal the signalling mediators that drive them.

## Introduction

The tumour microenvironment comprises a diverse array of cell types, including tumour cells, immune cells, fibroblasts, endothelial cells, and cancer-associated stromal cells, that interact dynamically to influence tumour progression, metastasis, and therapeutic response^1^. Whilst a range of approaches are commonly used to study these interactions, including transcriptomic profiling^2^, proteomics^3^, and flow cytometry^4^, these methods typically provide static snapshots of dynamic processes and require cell fixation or dissociation, which can disrupt native cellular behaviour. In contrast, live-cell imaging, particularly time-lapse microscopy, offers direct visualisation of cell co-cultures, capturing real-time cellular dynamics and interactions and thus fully preserving both spatial and temporal context.

One of the primary barriers to using time-lapse microscopy for studying cellular behaviour in co-culture systems is the analytical complexity of the resulting data, particularly due to challenges in tracking and segmenting heterogeneous, interacting cell populations^5^, as well as accurate cell type classification to enable cell-type-specific analyses of behaviour over time^6^. A common approach to determine the ground truth identity of cell types in microscopy images is to label them using fluorescent biomarkers. However, the combination of fluorescent staining and frequent light exposure during time-lapse imaging can disrupt normal cell behavior or induce phototoxicity, limiting its applicability in long-term experiments^7,8^. This motivates the use of label-free approaches for time-lapse microscopy, such as ptychography or phase-contrast microscopy, which enable for extended imaging over longer timescales without compromising cell viability or behaviour^9^. As a result, label-free imaging can generate substantially larger data sets, capturing single-cell dynamics over time. Effectively analysing such high-dimensional, time-resolved data requires advanced computational methods, including single-cell segmentation and tracking, feature extraction, and machine learning techniques capable of extracting meaningful biological insights from complex temporal patterns.

As a solution to this, we developed CellPhe^10^, a toolkit for automated cell phenotyping from microscopy timelapses, designed to extract and interpret quantitative features from complex cellular behaviours over time. Our previous work has showcased CellPhe’s utility in supporting biological research, including the characterisation of phenotypic heterogeneity in breast cancer cell responses to chemotherapy^10^, and multi-class phenotypic profiling across a panel of breast cancer cell lines^11^. In this work, we aim to demonstrate that CellPhe is also a valuable tool for the analysis of cell co-cultures, enabling detailed phenotypic characterisation of cellular interactions. By capturing dynamic behavioural changes at the single-cell level, CellPhe facilitates identification of interaction-driven phenotypes that may otherwise be missed through conventional population-level or static timepoint analyses.

As a proof-of-principle, here we apply the CellPhe toolkit to investigate phenotypic changes in breast cancer cells induced by co-culture with cancer-associated fibroblasts (CAFs). The presence of CAFs in the breast cancer tumour microenvironment has been widely reported to correlate with poor clinical prognosis^12,13^, as well as enhanced local invasion and distant dissemination of cancer cells^14^. Furthermore, CAFs have been shown to actively promote epithelial-to-mesenchymal transition (EMT) in breast cancer cells, leading to the loss of epithelial characteristics, increased migration and acquisition of mesenchymal traits^15^. This transition contributes to a more invasive and therapy-resistance phenotype, driving tumour progression and recurrence^16^. Given that CAFs induce significant phenotypic changes in breast cancer cells, we hypothesised that CellPhe would be capable of quantifying these changes and facilitating their interpretation in terms of underlying biological mechanisms. This capability enables researchers to investigate how intercellular interactions influence cell state transitions in a high-throughput, label-free manner. By leveraging time-resolved features, CellPhe provides a powerful framework for uncovering subtle behavioural signatures that reflect dynamic changes in cellular phenotype, such as those driven by co-culture conditions, without the need for molecular labels or endpoint assays.

As the phenotypic changes induced in breast cancer cells by CAFs are increasingly attributed to complex crosstalk between tumour and stromal cells, mediated by dynamic signalling through secreted cytokines, chemokines, and growth factors, we also performed a Luminex immunoassay to profile key signalling molecules within our co-culture experiments. These included TGF□β, IL□6, IL□8, VEGF, and HGF, key signalling molecules that are either secreted by CAFs or known to play a role in processes such as EMT, cell migration, and therapeutic response in breast cancer^15,17,18^. By correlating these secreted factor profiles with phenotypic features extracted from time-lapse imaging, we present a multi-modal approach for investigating the impact of CAF-derived signals on breast cancer cell behaviour, providing insights at both molecular and cellular levels.

## Methods

### Cell culture

The MDA-MB-231 cells were a gift from M. Djamgoz, Imperial College, London. SkBr3 cells were a gift from J. Rae, University of Michigan and MCF-10A cells were a gift from N. Maitland, University of York. MDA-MB-468 and MCF-7 cells were from ATCC. dsRed cancer associated fibroblast (CAF) cells were a gift from V. Speirs, University of Aberdeen. CAFs were stably transduced with recombinant lentivirus and were then sorted using FACS to enrich red fluorescent cells, CAFs were also hTERT-immortalised as described previously^19^. The molecular identity of all cell lines was verified by short tandem repeat analysis^20^.

Authenticated cell stocks were stored in liquid nitrogen and thawed for use in experiments. Thawed cells were subcultured 4-5 times prior to discarding and thawing a new stock to ensure that the molecular identity of cells was retained throughout. All cell lines, except for MCF-10A, were cultured separately in Dulbecco’s Modified Eagle Medium (DMEM) supplemented with 5% fetal bovine serum (FBS) and 4 mM L-glutamine. MCF-10A cells were cultured in DMEM/F12 (Invitrogen) with 5% heat-inactivated horse serum, 0.5 µg/mL hydrocortisone, 20 ng/mL human EGF, 10 µg/mL insulin, and 100 ng/mL cholera toxin. To minimise imaging artefacts, FBS was filtered using a 0.22 µm syringe filter before use. Cells were incubated at 37°C in plastic filter-cap T-25 flasks and were split at a 1:6 ratio during passaging. No antibiotics were added to the cell culture medium. Cells were confirmed to be free of mycoplasma before use in experiments through routine DAPI testing at monthly intervals.

### Co-culture experiments

For all experiments involving co-culture of MDA-MB-231 or MCF-7 with CAF cells, cell types were monocultured until seeding, where they were seeded together in 24-well plates 24 hours prior to imaging. In cases where the concentration of CAFs was increased, cells were seeded as cancer cell to CAF ratios of 3:1 (25% CAFs), 1:1 (50% CAFs) and 1:3 (75% CAFs), where the number of seeded cells/well remained consistent at 8000.

### CAF conditioned medium (CAF-CM)

CAFs were cultured in 25cm^2^ flasks until confluent and then their culture medium (referred to as CAF-CM throughout) was collected prior to sub-culturing. MDA-MB-231 or MCF-7 cells already seeded into 24-well plates then received a media change where either the collected CAF-CM was added or standard DMEM as a control. Cells were incubated in this media for 24 hours prior to imaging.

### Luminex experiments

#### Sample preparation

The MILLIPLEX® Human Circulating Cancer Biomarker Magnetic Bead Panel 1 (HCCBP1MAG-58K) and TGFβ1 Single Plex Magnetic Bead Kit (TGFBMAG-64K-01) were used to prepare medium samples for the Luminex immunoassay. It was necessary to seed samples into two separate 96-well plates prior to preparation for the Luminex due to the TGFβ1 single plex requiring an acid pre-treatment that is incompatible with the multi-plex panel. Tested samples included conditioned medium collected from MDA-MB-231, MCF-7 and CAF cells as well as from MDA-MB-231 and MCF-7 cells that had been cultured in CAF-CM for 72 hours prior to sample collection. Experimental variation was handled by testing 5 replicate wells for each sample. Biological variation was handled through replicate experiments, with samples collected from three separate experiments carried out on different weeks. Medium samples were centrifuged prior to seeding to ensure complete removal of cells or debris that would interfere with assay results. DMEM was added to background, standard and control wells to control for variation in analyte production as a result of sample culture medium. Magnetic beads for all analytes were mixed together in a mixing bottle provided within the kit and 25μl of this solution was added to all background, standard, control and sample wells. Plates were placed on a plate shaker and left to incubate overnight at 4°C. The following day, all wells were washed three times in the provided wash buffer following the plate washing instructions in the HCCBP1MAG-58K protocol and 25μl detection antibodies added to each well. Plates were then wrapped in foil and placed on a plate shaker at room temperature to incubate for 1 hour. 25μl of Streptavidin-Phycoerythrin was added to each well and plates re-wrapped in foil and placed on the plate shaker for a further 30 minute incubation. Well contents was then removed and plates washed a further three times prior to being read on the Luminex 200 system.

#### Data exportation and analysis

Data exported from the Luminex xPonent software was analysed in R using the `drLumi` and `limma` packages^21,22^. Analysis involved use of median fluorescent intensity (MFI) values and expected concentrations from standard wells for the fitting of standard curves. A 5-parameter log-logistic function was used for standard curve fitting, or a 4-parameter log-logistic function in cases where 5-parameter fitting did not converge. Models were then used for the prediction of analyte concentrations from raw MFI values for all samples. For any sample where concentration was predicted to be lower than the limit of quantification, the concentration for this sample was instead set as the lower limit.

#### Analyte gene mapping and EMT cross-referencing

Gene symbols corresponding to the protein analytes included in the MILLIPLEX® Human Circulating Cancer Biomarker Magnetic Bead Panel 1 panel were identified using GeneCards: The Human Gene Database (genecards.org). Analytes exhibiting statistically significant changes in concentration in the conditioned medium of MCF-7 or MDA-MB-231 cells following treatment with CAF-CM were selected for further analysis. These analytes were cross-referenced against two independent EMT gene datasets: msigdb M5930 Hallmark Epithelial Mesenchymal Transition (gsea-msigdb.org) and EMTome (emtome.org), both of which contain curated lists of genes associated with EMT processes.

### Image acquisition and exportation

On the day of imaging, cells were placed onto the Phasefocus Livecyte 2 (Phasefocus Limited, Sheffield, UK) to incubate for 30 minutes prior to image acquisition to allow for temperature equilibration. One 500μm x 500μm field of view per well was imaged to capture as many cells, and therefore data observations, as possible. Selected wells were imaged in parallel for 48 hours at 20x magnification with 6 minute intervals between frames, resulting in full time-lapses of 481 frames per imaged well. In cases where fluorescence was used, phase and fluorescence images were acquired in parallel for each well. Phasefocus’ Cell Analysis Toolbox® software was used for image processing, including background noise reduction with the rolling ball algorithm, as well as cell segmentation and tracking.

### Data analyses

#### CellPhe analyses

All phenotypic characterisation of cells was performed using the CellPhe toolkit (Python package, version 0.4.2). Within CellPhe, segmentation was performed with the integrated CellPose (cyto3 model) implementation, and tracking with the integrated TrackMate implementation (SparseLAP algorithm). Feature extraction was carried out using the `cell_features` and `time_series_features` functions, and optimal separation thresholds were determined using the `calculate_separation_scores` and `optimal_separation_features functions`. The full list of features extracted through CellPhe is provided in Wiggins et al. (2023)^10^.

#### Classification and clustering

Cell type classification was carried out using the classify_cells function from the CellPhe Python package (v0.4.2), which implements an XGBoost-based supervised classification framework. Separate binary classifiers were trained for MDA-MB-231 vs. CAF (training set: n = 814 MDA-MB-231, n = 680 CAF) and MCF-7 vs. CAF (training set: n = 1121 MCF-7, n = 557 CAF). Model performance was evaluated on independent test sets (n = 557 MDA-MB-231, n = 255 CAF; n = 557 MDA-MB-231, n = 255 CAF) and reported using accuracy and confusion matrices. Principal component analysis (PCA) was applied to z-scored feature data using scikit-learn. Independent test sets and co-culture datasets were projected into the PCA space derived from monoculture training data, enabling comparison of phenotypic profiles and visualisation of population-level shifts. To account for domain shift observed in co-culture experiments, unsupervised clustering was performed in PCA-reduced space using k-means (k = 2). Cluster identities were assigned based on their proximity to monoculture centroids, and validated against dsRed CAF fluorescence intensity as ground truth.

### Statistical analyses

All statistical analyses were carried out in Python (v3.12.3) using the SciPy, NumPy, and statsmodels libraries unless otherwise stated. For comparisons of phenotypic features between conditions, nonparametric Kruskal–Wallis (KW) tests were applied, followed by post hoc pairwise tests with False Discovery Rate (FDR) correction to account for multiple comparisons. To identify monotonic trends in phenotypic features across increasing CAF densities, Spearman’s rank correlation coefficients (ρ) were computed; features with |ρ| ≥ 0.2 and p < 0.05 were considered to show consistent directional change. For CAF-CM experiments, two-sample t-tests were used to assess differences in feature distributions between control and CAF-CM–treated cells, and results were visualised using volcano plots of log_2_ fold change versus –log_10_ p-value. For fluorescence intensity comparisons between classification- and clustering-based approaches, mean values ± standard error of the mean (SEM) were reported, and differences were interpreted in the context of CAF dsRed signal as a ground truth validation measure. In all cases, statistical significance thresholds were set at p ≤ 0.05 unless otherwise specified.

Statistical significance is denoted in figures as: p > 0.05 (n.s), p ≤ 0.05 (*), p ≤ 0.01 (**), and p ≤ 0.001 (***), p ≤ 0.001 (****).

## Results

CellPhe is an open-source toolkit for automated single-cell phenotyping from microscopy time-lapse data. Previous studies have demonstrated its utility in supporting biological research, including the characterisation of phenotypic heterogeneity in breast cancer cell responses to chemotherapy^10^, and multi-class phenotypic profiling across a panel of breast cancer cell lines^11^. The aim of this study was to assess whether CellPhe can extend beyond monoculture applications and enable accurate phenotypic analysis of mixed co-cultures of cells, allowing quantification of cell-cell interactions without the need for fluorescent labelling. As such, the experimental design consisted of two co-culture setups, in which CAFs were cultured with either MDA-MB-231 or MCF-7 breast cancer cells. ds-Red-labelled CAFs were used to provide ground truth information on true cell identity throughout the study, however, only label-free phenotypic features were used for all characterisation and classification analyses. The subsequent results focus on identifying phenotypic changes in MDA-MB-231 and MCF-7 cells induced by co-culture with CAFs, with the goal of elucidating potential interaction mechanisms and associated signalling pathways.

### Phenotypic characterisation of MDA-MB-231, MCF-7 and CAFs from microscopy time-lapse videos

To assess whether the phenotypic features extracted using CellPhe are sufficient to characterise breast cancer cell subtypes, time-lapse ptychography images of the breast cancer cell lines MDA-MB-231 and MCF-7, as well as CAFs, were acquired for each in monoculture, and time series features were extracted in preparation for supervised classification. Representative time-lapse images at 0h, 24h and 48h are provided in **Figure 1a**. CellPhe’s feature selection approach was then applied to reduce the initial set of 1,111 phenotypic time-series features to those most discriminative for two classification tasks: MDA-MB-231 vs. CAF and MCF-7 vs. CAF. This method calculates a measure of separation for each feature, defined as the ratio of between-groups variance to within-groups variance, and applies elbow-based thresholding to identify the most informative features. For the MDA-MB-231 vs. CAF comparison, 159 features exceeded the optimal separation threshold of 0.4, while 153 features exceeded an optimal threshold of 0.75 for the MCF-7 vs. CAF comparison. The features are categorised into texture, shape, size, and local cell density, with the following feature proportions: For MDA-MB-231 vs. CAF, texture accounted for 77%, shape for 10%, density for 9%, and size for 4%. For MCF-7 vs. CAF, texture accounted for 65%, shape 21%, density 10%, and size 5% **(Supplementary Figure 1, Supplementary Table 1 and 2)**. The refined list of features was used as input for principal component analysis (PCA), which identified distinct cell populations along PC1 for both MDA-MB-231 vs. CAF and MCF-7 vs. CAF. The variance explained by PC1 and PC2 was 52% and 11% for MDA-MB-231 vs. CAF, and 67% and 8% for MCF-7 vs. CAF, respectively **(Figure 1b)**. Independent test sets were projected onto the PCA space to assess whether the identified groupings were preserved in new data (n = 557 for MDA-MB-231, 1121 for MCF-7, and 255 for CAF). The projected points consistently aligned with the correct cell type clusters, demonstrating the generalisability and robustness of these phenotypic features in distinguishing cell populations **(Figure 1b)**.

**Figure 1.**
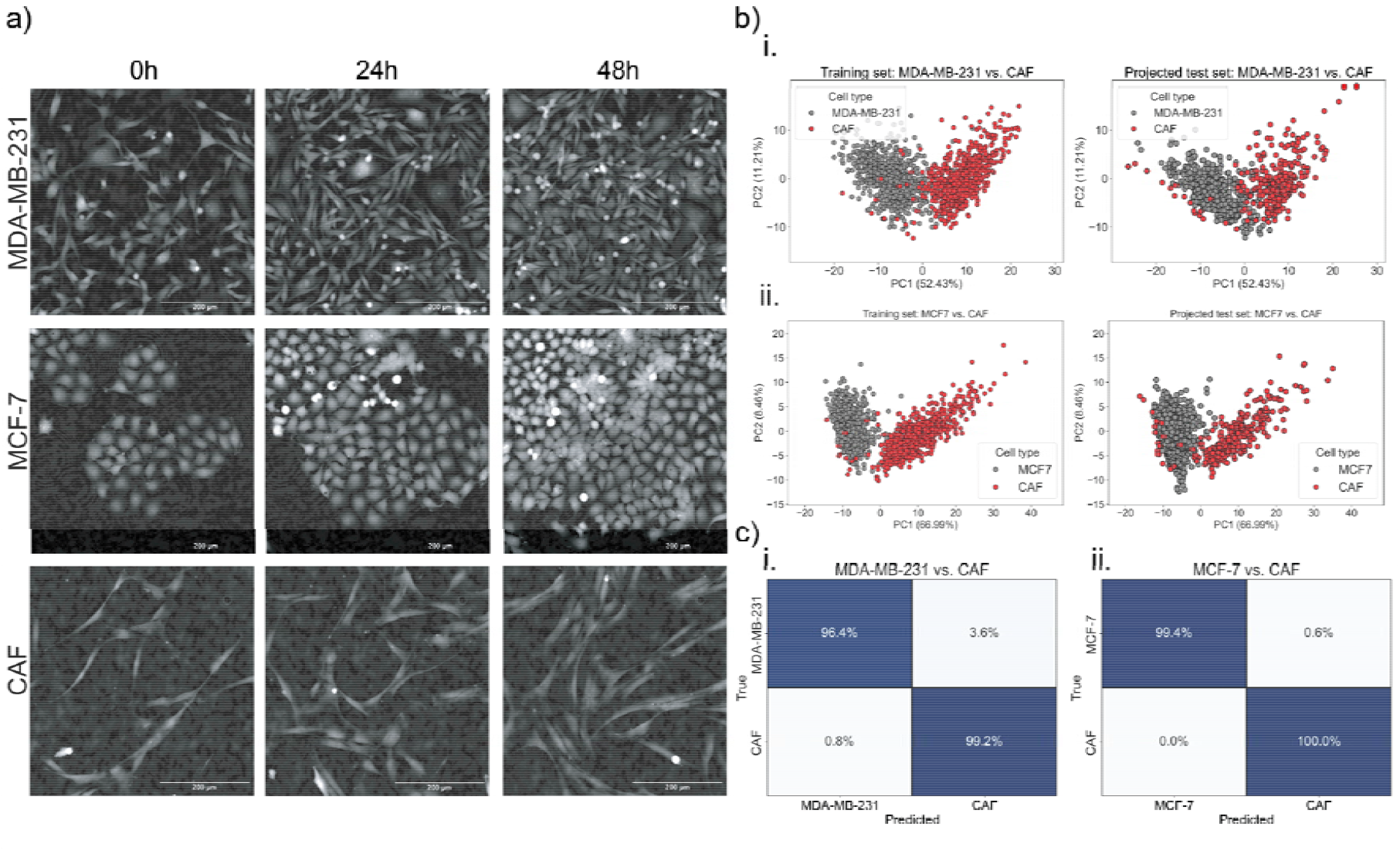
CellPhe enables phenotypic characterisation and classification of MDA-MB-231, MCF-7 and CAFs. **a)** Representative ptychographic images of MDA-MB-231, MCF-7 and CAF cells, demonstrating observable differences in their phenotypes. **b)** PCA scores plots for i. MDA-MB-231 vs. CAFs and ii. MCF-7 vs. CAFs. Left panels show training data colour-coded by true class labels, while right panels show independent test sets projected into the same PCA space, confirming the preservation of phenotypic separation. **c)** Confusion matrices for XGBoost classification of i. MDA-MB-231 vs. CAF (537/557 and 253/255 correctly classified) and ii. MCF-7 vs. CAF (1114/1121 and 255/255 correctly classified).

CellPhe’s XGBoost-based classification framework was used to train models capable of distinguishing between MDA-MB-231 and CAF, as well as between MCF-7 and CAF. When applied to independent test sets, the MDA-MB-231 vs. CAF model achieved classification accuracies of 96% for MDA-MB-231 (537/557 classified as MDA-MB-231) and 99% for CAF (253/255 classified as CAF), while the MCF-7 vs. CAF model achieved accuracies of 99% for MCF-7 (1114/1121 classified as MCF-7) and 100% for CAF **(Figure 1c)**.

### MDA-MB-231, MCF-7 and CAF classification from time-lapse videos of co-cultures

The successful performance of the MDA-MB-231 vs. CAF and MCF-7 vs. CAF classifiers suggested they would provide a useful tool for facilitating cell type identification from time-lapse images of co-cultures. Two sets of co-culture time-lapse experiments were conducted, one containing 50% MDA-MB-231 and 50% CAFs, and another containing 50% MCF-7 and 50% CAFs. For co-culture experiments, dsRed-labelled CAFs were used, and ptychography and red fluorescence images captured in parallel to provide ground truth labels for model validation. Representative phase and fluorescence images of the co-cultures are provided in **Figure 2a**, which demonstrate visibility of all cells within the ptychography channel but only CAFs in the red fluorescent channel.

**Figure 2.**
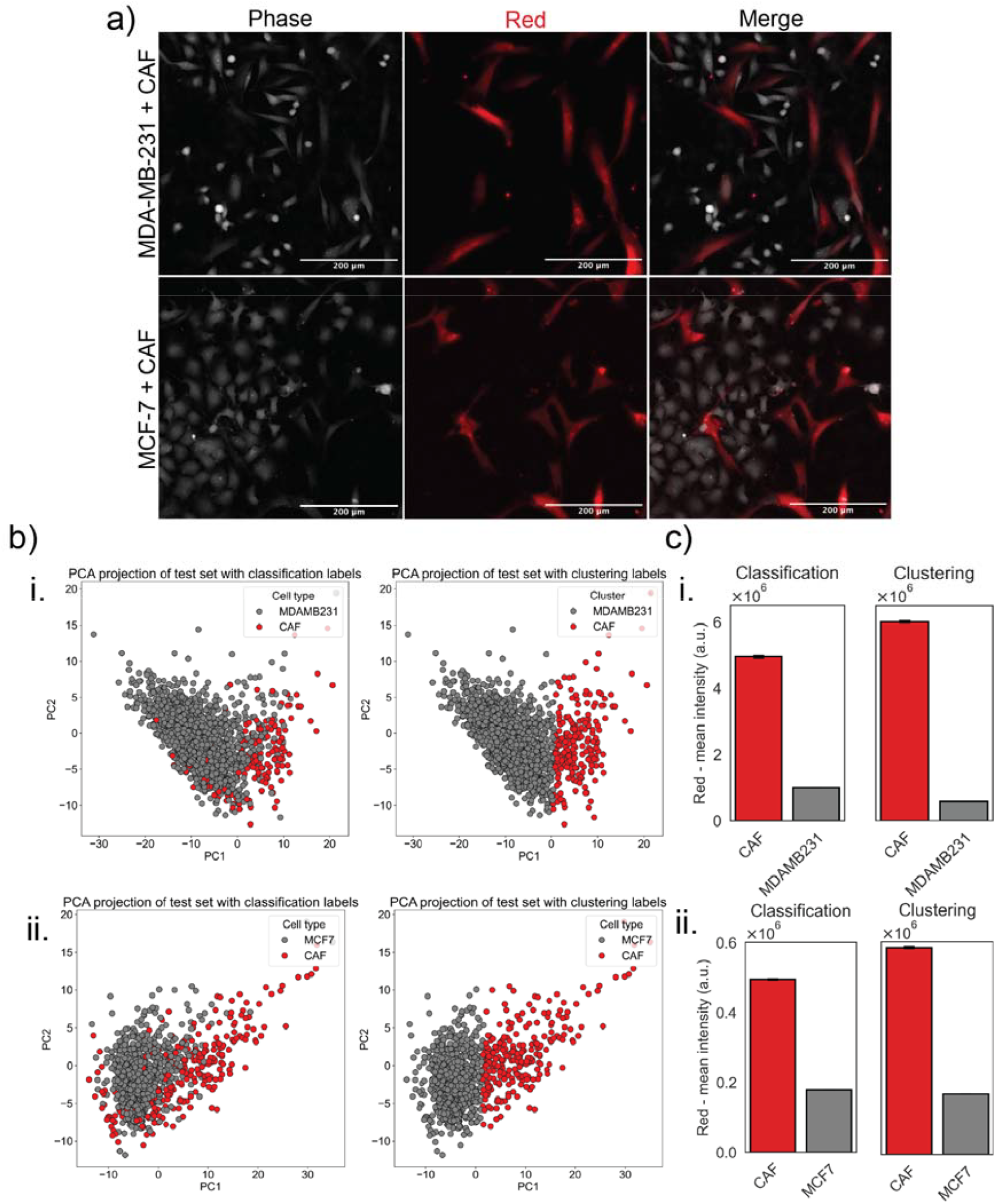
Co-culturing MDA-MB-231 and MCF-7 cells with CAFs induces a phenotypic domain shift. **a)** Representative images of MDA-MB-231 and MCF-7 cells co-cultured with dsRed CAFs, where ptychographic and fluorescence images were acquired in parallel. **b)** Co-culture data projected onto the monoculture PCA scores plot from Figure 1a for i. MDA-MB-231 and CAF, and ii. MCF-7 and CAF. Left panel shows predicted labels using the monoculture classifier, while the right panel shows labels obtained via k-means clustering and centroid pairing. Points are colour-coded according to predicted class labels. **c)** Bar plots of mean red fluorescent intensity of tracked cells for i. MDA-MB-231 vs. CAF and ii. MCF-7 vs. CAF. Cells are grouped by predicted class label, with the monoculture classifier used for the plots on the left (labelled “Classification”) and k-means clustering with centroid pairing on the right (labelled “Clustering”). The plots demonstrate improved cell type identification using the clustering approach, with increased mean fluorescent intensity for CAFs (4.95×10□ ± 3.78×10□ to 6.01×10□ ± 3.53×10□ for MDA-MB-231 vs. CAF and 4.95×10□ ± 2.87×10^3^ to 5.85×10□ ± 3.19×10^3^ for MCF-7 vs. CAF) and decreased mean fluorescent intensity for MDA-MB-231 (9.99×10□ ± 7.77×10^3^ to 5.75×10□ ± 4.76×10^3^) and MCF-7 (1.79×10□ ± 5.15×10^2^ to 1.71×10□ ± 4.50×10^2^). Barplots show mean ± SEM.

Projection of co-culture data onto the PCA space from **Figure 1b** demonstrated a loss of clearly defined clusters and a shift in PC1 and PC2 centroids for all cell types, indicating a phenotypic shift likely driven by interactions between CAFs and cancer cells (**Figure 2b**). This domain shift suggests that cell behaviour and morphology are altered in co-culture conditions compared to monoculture. As a result, classifier performance is impacted, with unclear groupings of cancer cells and CAFs observed following model inference (**Figure 2b**). To address poor classification performance as a result of domain shift, k-means clustering was applied to identify two distinct clusters within the PCA space, with each cluster being labelled based on its proximity to the nearest centroid from the monoculture PCA space. As only CAFs express red fluorescence, we used red channel intensity to validate cell identities predicted by both classification and clustering approaches. The clustering-based method yielded higher mean red fluorescence in the predicted CAF group and lower intensities in MDA-MB-231 and MCF-7 cells, indicating improved prediction of cell types and fewer misclassifications when using the clustering-based approach (**Figure 2c**).

The intensity for predicted CAFs increased from 4.95×10□ ± 3.78×10□ to 6.01×10□ ± 3.53×10□ in the MDA-MB-231 vs. CAF labelling, and from 4.95×10□ ± 2.87×10^3^ to 5.85×10□ ± 3.19×10^3^ in the MCF-7 vs. CAF labelling. In contrast, fluorescent intensity for the predicted MDA-MB-231 and MCF-7 groups decreased, from 9.99 × 10□ ± 7.77 × 10^3^ to 5.75 × 10□ ± 4.76×10^3^ for MDA-MB-231 cells, and from 1.79×10□ ± 5.15×10^2^ to 1.71×10□ ± 4.50×10^2^ for MCF-7 cells. Examples of cells classified for each cell type are shown in **Supplementary Figure 2**.

### CAFs modulate MDA-MB-231 and MCF-7 phenotypes in a density-dependent manner

Having identified a phenotypic domain shift in the co-culture datasets, we next aimed to pinpoint the specific phenotypic features of MDA-MB-231 and MCF-7 cells that were altered by CAFs, by directly comparing their behaviour in monoculture vs. co-culture. MDA-MB-231 and MCF-7 cells were co-cultured with increasing densities of CAFs, including 0%, 25% (3:1 ratio), 50% (1:1 ratio), and 75% (1:3 ratio) CAFs. Representative phase and red fluorescence images are provided in **Figure 3a**. Identification of cells using the k-means and centroid pairing approach showed an expected increase in the proportion of predicted CAFs with higher seeding densities. In co-culture with MDA-MB-231 cells, the proportion of predicted CAFs increased from 6% at 0% CAFs to 14%, 40%, and 66% at 25%, 50%, and 75% CAFs, respectively. Similarly, in co-culture with MCF-7 cells, predicted CAFs rose from 5% at 0% CAFs to 30%, 56%, and 87% with increasing CAF densities (**Supplementary Figure 3**). Cells identified as MDA-MB-231 and MCF-7 through this approach were then used for the subsequent analysis.

**Figure 3.**
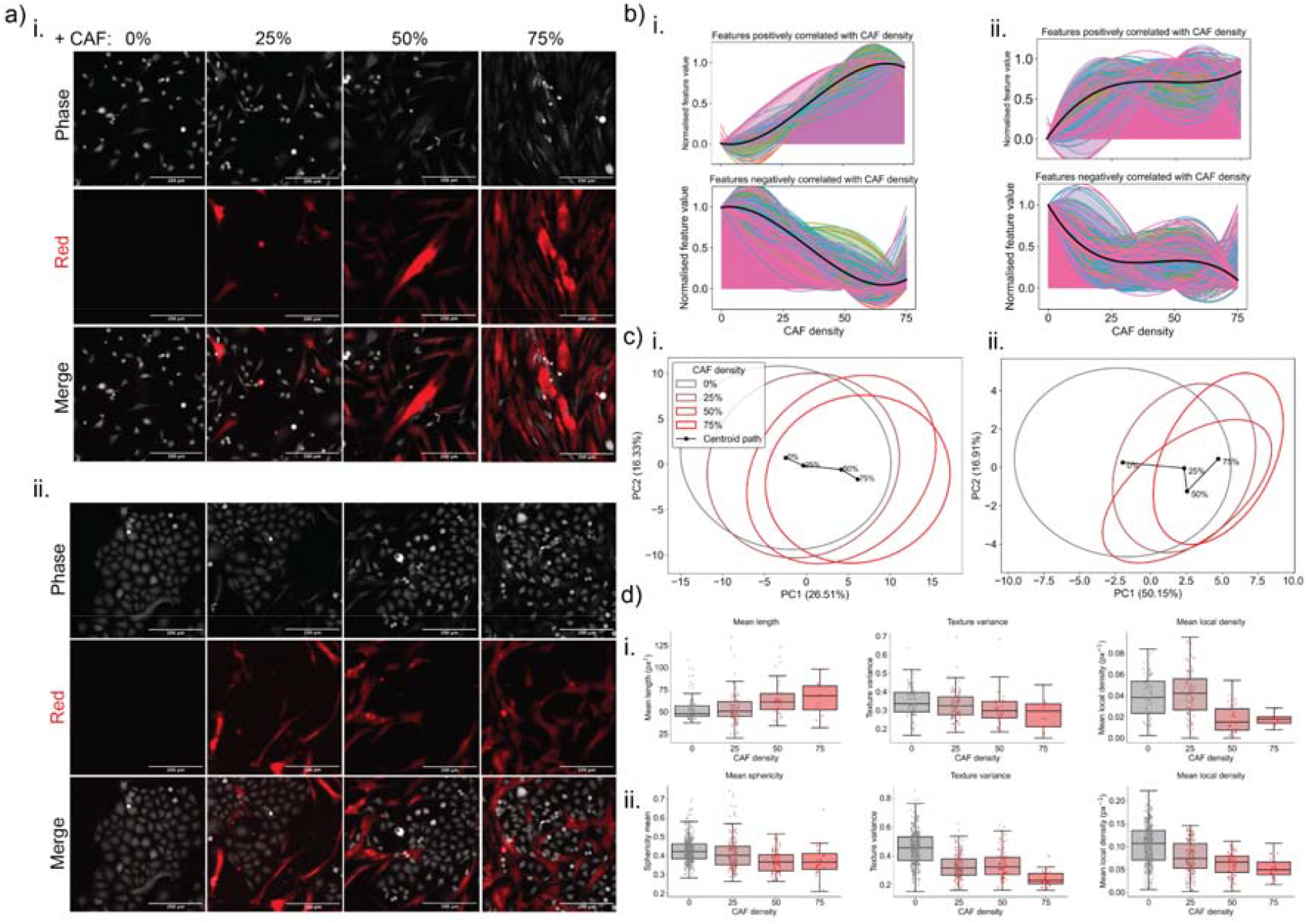
Co-culture with CAFs induces concentration-dependent phenotypic changes in MDA-MB-231 and MCF-7 cells. **a)** Representative ptychographic and red fluorescence images for dsRed CAFs co-cultured with i. MDA-MB-231 and ii. MCF-7. Note that 0%, 25%, 50% and 75% CAFs refer to cancer cell to CAF seeding ratios of 1:0, 3:1, 1:1 and 1:3 respectively. **b**) Area plots show normalised mean values of features that exhibited directional trends (absolute Spearman’s ρ ≥ 0.2, p□<□0.05) across increasing CAF concentrations (0%, 25%, 50%, 75%). Features were separated into those that i. increased or ii. decreased with CAF concentration. Each coloured area represents one feature, and the bold black line indicates the average trend across all positively or negatively correlated features. **c**) PCA score plots for i. MDA-MB-231 and ii. MCF-7 based on features shown to significantly vary with increasing CAF density, as determined by Kruskal-Wallis and Spearman’s rank correlation. PCA scores plots show phenotypic divergence of cancer cells with increasing CAF concentration (0%, 25%, 50%, 75%). 95% confidence ellipses are shown for each concentration, with the black centroid path highlighting progressive separation from the control (0%) as CAF concentration increases, indicating dose-dependent shifts in cancer cell phenotype. **d)** Boxplots of example features that showed positive or negative correlations with increasing CAF density for (i) MDA-MB-231 and (ii) MCF-7 cells. Shown are an increase in mean length for MDA-MB-231 cells and a decrease in sphericity for MCF-7 cells, as well as decrease in texture variance and local cell density for both cell types.

We applied the Kruskal–Wallis nonparametric test to each CellPhe-derived feature, with False Discovery Rate (FDR) correction to control for multiple comparisons, in order to identify phenotypic changes in breast cancer cells across increasing CAF densities. This analysis revealed 358 significantly altered features in MDA-MB-231 cells and 923 in MCF-7 cells, highlighting substantial phenotypic shifts in both lines. To determine the direction of these changes, we next used Spearman’s rank correlation to assess monotonic trends with CAF density. Features with an absolute Spearman correlation coefficient greater than 0.2, a threshold chosen to capture modest but consistent responses, were classified as increasing or decreasing. In MDA-MB-231 cells, 62 features increased and 135 decreased with rising CAF density, while in MCF-7 cells, 148 features increased and 247 decreased. Together, these findings demonstrate that both breast cancer cell lines undergo marked phenotypic adaptations in response to co-culture with CAFs (**Figure 3b, Supplementary Table 3 and 4**).

The subset of features that passed both the Kruskal–Wallis (KW) significance test and the Spearman trend filter were then used as input for PCA. This analysis revealed that as CAF density increased, the phenotypes of both MDA-MB-231 and MCF-7 cells progressively diverged from their monoculture states (**Figure 3c**). The PCA scores plot illustrates this shift, with 95% confidence ellipses showing reduced overlap with, and increased traversal away from, the monoculture ellipse as CAF density increases. In both cell lines, the primary centroid migration occurred along PC1, which explained 27% of the variance in MDA-MB-231 cells and 50% in MCF-7 cells. This result indicates that CAF density–driven phenotypic variation is largely captured along the first principal component.

Directional feature changes highlighted alterations in cell shape, although different descriptors were more informative for each cell line. The mean length of MDA-MB-231 cells increased from 53.2 ± 14.8 µm in monoculture to 64.6 ± 19.3 µm at 75% CAFs. While the KW test indicated significant overall differences (H = 15.77, p = 0.0013), post hoc testing showed that increased elongation was most evident at 50% CAFs **(Figure 3d**).

In contrast, for MCF-7 cells, which are typically more rounded and compact, length was less descriptive; instead, changes in sphericity captured their morphological response. Sphericity decreased significantly in MCF-7 cells with increasing CAF density, from 0.428 ± 0.067 in monoculture to 0.370 ± 0.088 at 75% CAFs (KW, H = 57.7, p < 0.0001). Post hoc testing confirmed significant reductions at 25% (p = 0.008), 50% (p < 0.0001), and 75% CAFs (p = 0.0004) compared to monoculture. These results indicate that MCF-7 cells progressively lose their rounded epithelial morphology and adopt a less spherical, more mesenchymal-like shape under CAF influence.

In ptychographic imaging, pixel intensity reflects cellular dry mass, allowing texture features to capture how intracellular material is organised. One such feature, Cooc01Var_asc, which measures the temporal ascent of pixel intensity variance, decreased significantly with increasing CAF density in both breast cancer cell lines. In MDA-MB-231 cells, Cooc01Var_asc decreased from 0.348 ± 0.082 in monoculture to 0.278 ± 0.084 at 75% CAFs (KW, H = 13.327, p = 0.004), while in MCF-7 cells it declined from 0.453 ± 0.116 to 0.242 ± 0.054 (KW, H = 198.874, p < 0.0001). This reduction indicates that fluctuations in dry mass distribution became less pronounced over time, consistent with a more stable intracellular organisation. Such stabilisation is characteristic of a mesenchymal-like state, in which cells adopt a more polarised and persistent morphology that supports migration.

Furthermore, local cell density, measured as the number of cells within a defined spatial radius around each cell, exhibits a decreasing trend in both MDA-MB-231 (from 0.040 ± 0.019 in monoculture to 0.018 ± 0.006 at 75% CAFs, KW, H = 47.379, p < 0.0001) and MCF-7 cells (from 0.105 ± 0.042 in monoculture to 0.052 ± 0.022 at 75% CAFs, KW, H = 119.540, p < 0.0001) as the proportion of CAFs increases. This decrease in MDA-MB-231 cells can be attributed to their alignment with CAFs (**Supplementary Figure 4**), which occupy a larger area, thereby decreasing the number of neighbouring cells that fall within the defined measurement radius used for density calculation. In the case of MCF-7 cells, this reduction in local cell density reflects a loss of their typical epithelial clustering behaviour, with cells dispersing and adopting a more scattered arrangement in response to CAF co-culture.

These features are hallmarks of a mesenchymal phenotype, characterised by elongated morphology, loss of epithelial cell–cell adhesion, reduced local density, and stabilisation of intracellular organisation. Together, they indicate a shift toward a polarised state typically associated with migratory capacity. In co-culture, however, these morphological and organisational changes were not paralleled by significant alterations in motility, suggesting that direct CAF contact may constrain increased cancer cell movement even as it promotes mesenchymal-like traits.

### Culturing with CAF-conditioned medium induces changes to MDA-MB-231 and MCF-7 phenotypes

To assess whether phenotypic changes in cancer cells could be driven by CAF-secreted or - depleted factors, such as chemokines and cytokines, rather than by direct cell-cell contact, MDA-MB-231 and MCF-7 cells were cultured in CAF-conditioned medium (CAF-CM), and compared to MDA-MB-231 and MCF-7 cells grown in standard culture medium.

Representative time-lapse images are shown in **Figure 4a**. Based on visual inspection, MDA-MB-231 cells cultured in CAF-CM appeared more elongated, while MCF-7 cells displayed greater variation in shape, including elongated morphologies uncharacteristic of this line and more typical of mesenchymal-like cells. In addition, MCF-7 cells formed fewer colonies, with individual cells more often migrating away from one another. To quantify both the magnitude and statistical significance of CAF-CM–induced phenotypic changes, log_2_ fold changes and p-values from two-sample t-tests were calculated for all features extracted using CellPhe. These results are visualised using volcano plots in **Figure 4b**. The analysis showed that CAF-CM significantly altered the phenotypic profiles of both cell lines. For MDA-MB-231 cells, 787 out of 1,111 features (≈ 71%) exhibited significant changes (p < 0.01) upon culture with CAF-CM, while for MCF-7 cells, 566 out of 1,111 features (≈ 51%) were significantly altered.

**Figure 4.**
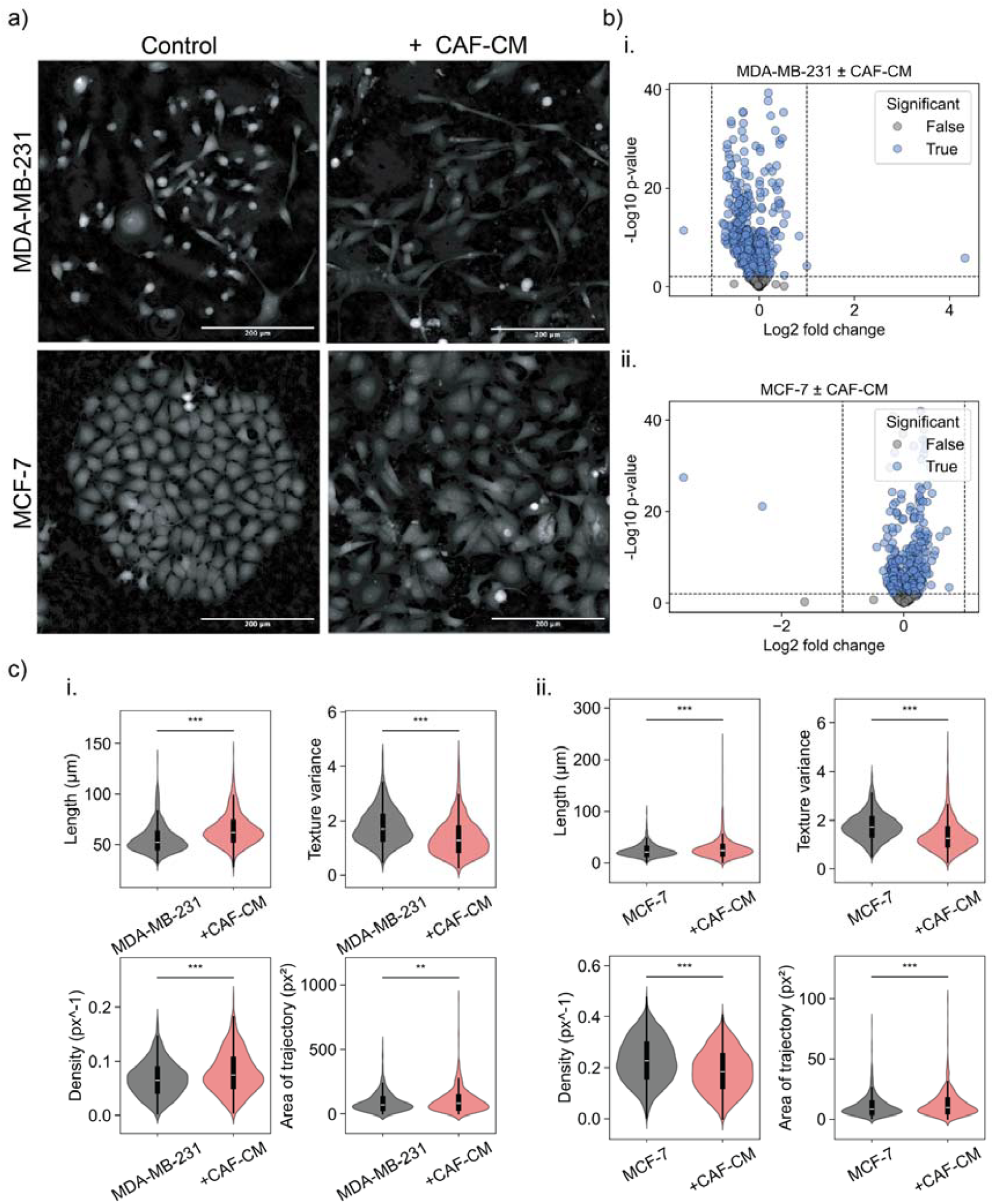
Co-culture with CAF-CM induces phenotypic changes in MDA-MB-231 and MCF-7 cells. **a)** Representative time-lapse images of MDA-MB-231 and MCF-7 cells cultured in standard culture medium or CAF-CM. **b)** Volcano plots depicting phenotypic changes induced by CAF-CM in i. MDA-MB-231 and ii. MCF-7 cells. Points are coloured blue for significant changes (p ≤ 0.01), and grey for non-significant changes. The dashed horizontal line at -log_10_ p-value = 2 indicates a threshold for statistical significance, while the dashed vertical lines at log_2_ fold change of ±1 denote a 2-fold change. **c)** Violin plots showing selected phenotypic features with significant changes induced by CAF-CM in i. MDA-MB-231 and ii. MCF-7 cells. Significant differences between control and CAF-CM conditions are indicated, with statistical significance determined by two-sample t-tests (** = p ≤□ 0.01, *** = p ≤□ 0.001).

Features that underwent significant changes following culture with CAF-CM were indicative of a shift toward more mesenchymal and aggressive phenotypes, consistent with those observed during direct co-culture with CAFs (**Figure 4c, Supplementary Table 5**). Notably, both MDA-MB-231 and MCF-7 cells exhibited increased elongation and reduced sphericity following exposure to CAF-CM. For example, the mean cell length increased from 56±14μm to 65±15μm in MDA-MB-231 cells (p ≤ 0.001), and from 55±8□μm to 57±12□μm in MCF-7 cells (p ≤ 0.001).

In co-culture, this stabilisation was reflected by reduced variance in pixel intensity distribution over time, whereas in CAF-CM it was captured through a reduction in fluctuations of overall cellular mass, with the total ascent in mean pixel intensity across the time series decreasing from 1.80 ± 0.65 to 1.40 ± 0.66 in MDA-MB-231 cells (p ≤ 0.001), and from 1.76 ± 0.55 to 1.38 ± 0.66 in MCF-7 cells (p ≤ 0.001). A reduction in fluctuations of average cellular mass over time indicates that cells are no longer continually reorganising their internal contents, but instead maintain a more fixed intracellular architecture. Such stabilisation is a recognised feature of EMT, where cells transition to a polarised, elongated state that supports persistent migration^23^.

Unlike in co-culture, where movement features remained unchanged, cancer cells grown in CAF-CM exhibited significantly greater migration. This was quantified using the trajectory area, which measures the two-dimensional space a cell covers during the timelapse, with larger trajectory areas indicating that cells travelled further from their starting position, consistent with increased motility. In MDA-MB-231 cells, the mean trajectory area increased from 91 ± 76 μm^2^ in monoculture to 104 ± 91 μm^2^ in CAF-CM (p < 0.01). Similarly, in MCF-7 cells, the trajectory area increased from 11 ± 9 μm^2^ to 13 ± 1 μm^2^ 1 (p < 0.001). The absence of a corresponding effect in co-culture may reflect the physical constraints imposed by CAFs themselves, which can hinder cancer cell displacement despite promoting mesenchymal-like morphology and intracellular stabilisation.

Decrease in local cell density for MCF-7 cells cultured in CAF-CM (from 0.23 ± 0.09 to 0.19 ± 0.08, p ≤□ 0.001), as was observed in co-culture data sets, again suggests a loss of polarity and the transition towards a more mesenchymal-like state associated with decreased cell-cell adhesion or altered migration patterns. In contrast, MDA-MB-231 cells experienced an increase in local cell density (0.068 ± 0.03px^−1^ to 0.079 ± 0.04px^−1^, p ≤□ 0.001). This observation is consistent with the timelapse videos of MDA-MB-231 cells cultured in CAF-CM, which showed a tendency for closely positioned cells to align and migrate collectively (**Supplementary Figure 5**). These findings suggest that CAF-CM not only promotes mesenchymal traits but may also facilitate coordinated movement and local clustering of MDA-MB-231 cells.

### Microenvironmental changes in MDA-MB-231 and MCF-7 cells cultured with CAF-CM

CAF-CM-induced phenotypic changes in MDA-MB-231 and MCF-7 cells, consistent with an increase in mesenchymal-like characteristics, led us to investigate whether secreted factors within CAF-CM are associated with EMT. A Luminex multiplex assay of 25 analytes was run to determine the presence or absence of certain soluble factors within cultured medium samples. The test panel consisted of magnetic beads for the analysis of AFP, Total PSA, CA15-3, CA19-9, MIF, TRAIL, Leptin, IL-6, sFasL, CEA, CA125, IL-8, HGF, sFas, TNFα, Prolactin, SCF, CYFRA 21-1, OPN, FGF2, β-HCG, HE4, TGFα, VEGF-A and TGF-β.

The heatmap of scaled MFI values (**Figure 5a**) showed the presence of two main analyte clusters within the dataset: one for analytes in high abundance within CAF-CM, and one for analytes with lower abundance. The high-abundance cluster included TGF-β, IL-8, OPN, sFas, IL-6, HGF, and VEGF-A. Across both MDA-MB-231 and MCF-7 cells, these factors were present at relatively low levels in controls but increased markedly in CAF-CM. For example, IL-6 rose from 330 ± 257 to 16,453 ± 4565 MFI in MDA-MB-231 cells (log□FC = 5.64, q = 0.059) and from 414 ± 622 to 14,739 ± 5999 MFI in MCF-7 cells (log□FC = 5.16, q = 0.095). Similarly, IL-8 increased from 3748 ± 2136 to 16,217 ± 878 MFI in MDA-MB-231 cells (log□FC = 2.11, q = 0.015) and from 1298 ± 1692 to 14,467 ± 451 MFI in MCF-7 cells (log□FC = 3.48, q = 0.013). TGF-β also showed consistent increases, from 1716 ± 669 to 6172 ± 706 MFI in MDA-MB-231 cells (log□FC = 1.85, q = 0.010) and from 1760 ± 840 to 5775 ± 555 MFI in MCF-7 cells (log□FC = 1.71, q = 0.013). Other analytes in this cluster, such as OPN, sFas, and HGF, showed large fold increases (e.g., OPN increased ∼36-fold in MDA-MB-231 and ∼13-fold in MCF-7), but owing to greater variability between replicates their changes did not consistently reach significance after FDR correction. VEGF-A exhibited only modest increases in both cell lines (MDA-MB-231: 2328 ± 3019 to 4364 ± 1862 MFI; MCF-7: 603 ± 408 to 1736 ± 732 MFI), which were also not significant after correction.

**Figure 5.**
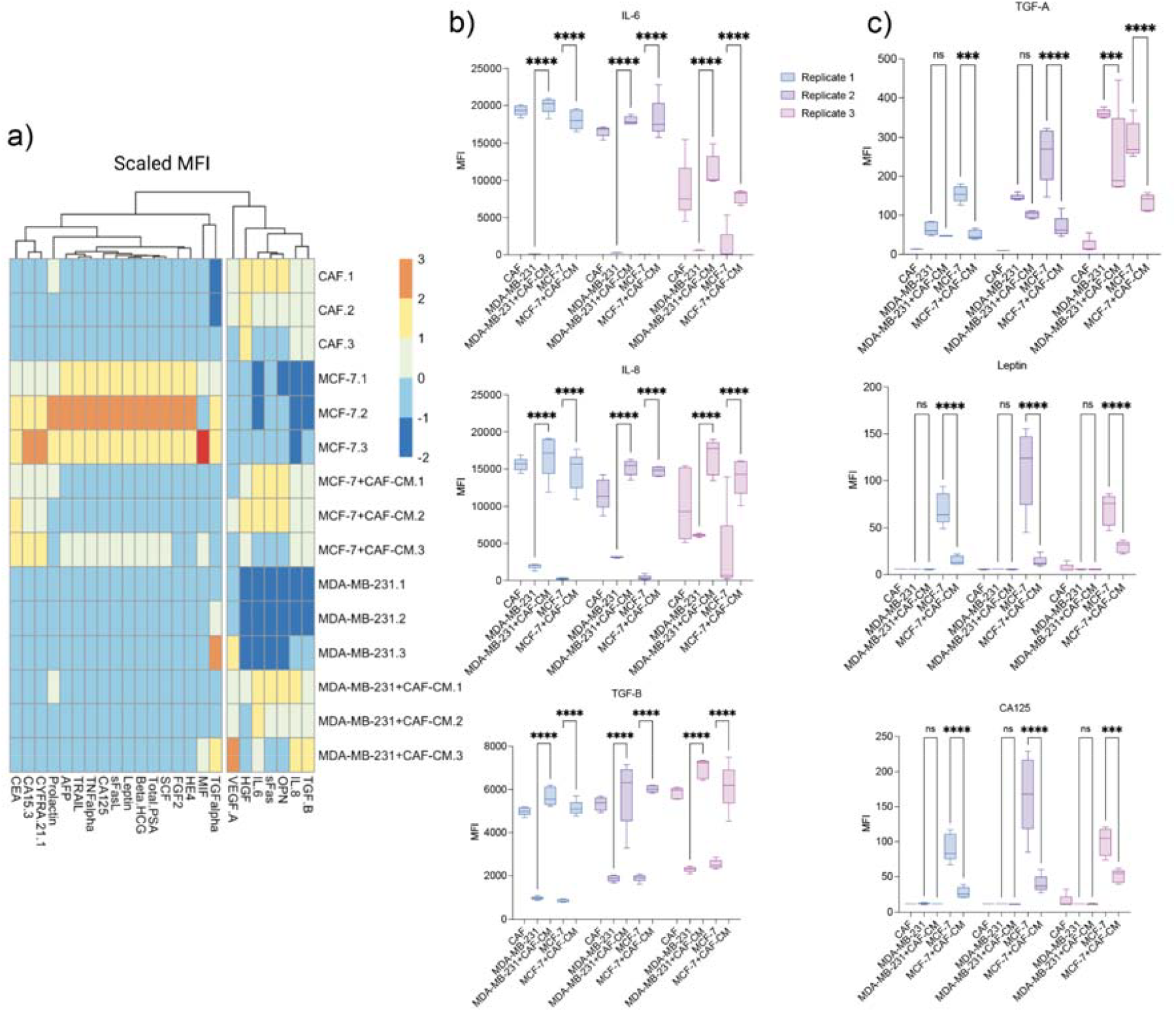
CAF-CM induces changes in analyte production in MDA-MB-231 and MCF-7 cells. **a)** Heatmap to aid visualisation of differences in analyte production of MDA-MB-231 and MCF-7 induced by culture with CAF-CM. Note that MFI values were first averaged to provide mean values for each of three experimental replicates, values were then normalised to allow changes on different scales to be compared. The heatmap demonstrates that addition of CAF-CM increases the presence of certain analytes within MDA-MB-231 and MCF-7 culture medium, such as TGF-B, IL-6 and OPN. The addition of CAF-CM reduces the presence of certain analytes within MCF-7 medium such as CA15-3, TGF-A and TNF-A. **b)** The addition of CAF-CM significantly increased the presence of IL-6, IL-8 and TGF-B in MDA-MB-231 and MCF-7 medium samples across all three experimental replicates, highlighting changes to cancer cell microenvironment induced by CAFs. **c)** The addition of CAF-CM significantly decreased the presence of TGF-A, leptin and CA-125 in MCF-7 cells across all three experimental replicates, suggesting a cross-talk between cancer cells and CAFs in which secretions from CAFs inhibit certain secretions from cancer cells.

In contrast, the second cluster comprised analytes abundant in the medium of MCF-7 cells, including TGFα, MIF, HE4, FGF2, SCF, Total PSA, β-HCG, Leptin, sFasL, CA125, TNFα, TRAIL, AFP, Prolactin, CYFRA 21-1, CA15-3, and CEA. Strikingly, the addition of CAF-CM led to reductions in most of these factors, particularly in MCF-7 cells. For example, HE4 decreased from 31 ± 3.8 to 14.6 ± 2.5 MFI (log_2_FC = –1.09, q = 0.072), FGF2 from 12.0 ± 1.1 to 8.9 ± 0.1 MFI (log_2_FC = –0.43, q = 0.093), Leptin from 84.3 ± 25.3 to 19.6 ± 8.8 MFI (log_2_FC = –2.10, q = 0.093), TRAIL from 245.9 ± 62.3 to 60.7 ± 32.2 MFI (log_2_FC = –2.02, q = 0.088), and AFP from 119.8 ± 24.3 to 46.0 ± 11.0 MFI (log_2_FC = –1.38, q = 0.088).

Prolactin was also reduced, from 165 ± 45.6 to 66.6 ± 27.9 MFI (log_2_FC = –1.31, q = 0.093). Several analytes, including MIF, CA125, CYFRA 21-1, and CEA, showed downward trends but did not reach significance, with CEA in particular showing little change (151 ± 65.5 to 154 ± 79.9 MFI, log_2_FC ≈ 0, q ≈ 0.97). In MDA-MB-231 cells, changes were less pronounced, although SCF increased (10.1 ± 1.9 to 21.2 ± 3.7 MFI, log_2_FC = 1.07, q = 0.14) while sFasL decreased (11.1 ± 1.0 to 8.3 ± 0.8 MFI, log_2_FC = –0.42, q = 0.14).

Barplots are provided in **Figure 5b** and **Figure 5c** to further aid visualisation of the changes to MFI for certain analytes induced by the addition of CAF-CM. Additionally, barplots of EMT-specific analytes identified by cross-referencing soluble factors showing significant changes in the culture medium after CAF-CM treatment to two independent EMT signature databases (msigdb and EMTome) are shown in **Supplementary Figure 6**. MFI values for IL-6, IL-8 and TGF-β are shown in **Figure 5b**. These plots show the presence of IL-6, IL-8 and TGF-β within CAF-CM, as well as a highly significant increase within MDA-MB-231 and MCF-7 samples where CAF-CM had been added, across all 3 experimental replicates. These plots suggest an additive effect in which addition of CAF-CM to MDA-MB-231 and MCF-7 cells results in a culture medium that contains the summation of secretions from each cell type. MFI values for TGFα, leptin and CA-125 are shown in **Figure 5c** and **Supplementary Figure 6**. The secretion of leptin and CA-125 from MCF-7 cells was significantly decreased following culture with CAF-CM, compared to MCF-7 cells cultured in normal medium for the same period, suggesting an inhibitory effect on such analytes mediated by factors secreted by CAFs. The same was observed for TGFα, with significant decrease in production by MCF-7 cells following culture with CAF-CM. Interestingly, no significant decrease in TGFα was observed for MDA-MB-231 cells following culture with CAF-CM for experimental replicates 1 and 2, although a significant decrease was observed in replicate 3 where the original MFI for TGFα within the control MDA-MB-231 sample was much higher than for the other replicates. This discrepancy could suggest that inhibition of TGFα in MDA-MB-231 cells by CAFs is dependent on the concentration of TGFα produced by the cancer cells, with greater concentrations resulting in a greater inhibitory response by CAFs. Whereas, the observed decrease in EMT-specific analytes in the medium of MCF-7 cells following CAF-CM treatment may suggest a context-dependent modulation of epithelial plasticity, and may reflect a more nuanced cell-specific response to stromal signalling.

In summary, the results of this study have shown how CellPhe’s phenotypic profiling is able to not only accurately distinguish breast cancer cell populations, but also reveal important CAF-driven, density-dependent EMT-like transitions, marked by elongation, intracellular reorganisation and increased migration, captured through time-lapse imaging. These shifts, which were replicated by CAF-CM and validated by Luminex experiments, underscore the secretory influence of CAFs in modulating cancer cell behavior and affirm CellPhe’s power in decoding dynamic phenomic signatures of malignancy.

## Discussion

Using microscopy to assess changes to cell behaviour induced by co-culturing with different cell types is currently a challenging task due to the complexities that arise when trying to separate out cell types following image acquisition. The results presented here show that our toolkit for automated cell phenotyping, CellPhe, is able to accurately classify cell types within co-cultures using their phenotypic characteristics in order to isolate populations for downstream analysis. Current approaches for isolation of specific cell populations include flow cytometry and fluorescence-activated cell sorting, though such approaches involve consideration of only a handful of pre-determined metrics for characterisation of cell types and have the potential to induce stress within cells^24^. In contrast, CellPhe facilitates accurate isolation of cell types using a rich panel of label-free features, enabling interpretable feature selection without negatively impacting cell health.

Culturing MDA-MB-231 and MCF-7 cells with CAFs as well as CAF-CM induced phenotypic changes in both cell lines, with cells displaying more elongated cell shape, decreased colony formation, and stabilisation of intracellular components. Together, these hallmarks are consistent with a shift towards a more aggressive, mesenchymal-like phenotype. These findings strongly suggest that CAFs drive EMT in both cancer cell lines, in line with previous reports of CAF-induced EMT in breast cancer^15^. Our findings reinforce the emerging view that CAFs act as central orchestrators of epithelial plasticity within the tumour microenvironment, shaping breast cancer cell behaviour through EMT induction and thereby promoting aggressive disease phenotypes. Importantly, we also demonstrate a new microscopy-based approach for identifying these dynamic changes directly from time-lapse data.

Our findings showed that the magnitude of these phenotypic shifts was modulated by CAF abundance, with higher CAF density driving progressively larger deviations from monoculture phenotypes. This observation aligns with previous findings that CAF-rich tumours are often more aggressive ^25–27^, and provides a mechanistic basis for how CAF density in the tumour microenvironment may scale EMT induction. Interestingly, while cells exhibited mesenchymal-like morphologies in direct co-culture, motility was not increased under these conditions, in contrast to CAF-CM where motility increased significantly. This difference suggests that CAF contact imposes physical constraints even as it promotes EMT-like states, whereas soluble signalling alone promotes cell dispersal and migration.

The Luminex immunoassay provided additional, complementary information to the phenotypic metrics extracted from time-lapse images, providing a list of biomarkers that may mediate the phenotypic changes induced in cancer cells by CAFs. The use of a pre-existing multiplex panel meant that biomarkers with limited associations to CAFs within the literature, such as CA-125, were also able to be assessed with no prior hypothesis of their involvement in CAF-cancer cell interactions. The Luminex assay showed the presence of analytes known to be secreted by CAFs within CAF-CM such as TGF-B, IL-6 and IL-8, with all of these analytes also present within medium collected following culturing of MDA-MB-231 and MCF-7 cells in CAF-CM. Such analytes were at relatively low concentrations within the control medium samples, demonstrating how the presence of CAF secretions alters the microenvironment of cancer cells and suggesting their potential role in inducing the observed phenotypic changes in cancer cells following culture with CAFs.

By cross-referencing analytes significantly altered by CAF-CM treatment with curated EMT signature databases (MSigDB and EMTome), we identified a subset of factors with strong associations to EMT regulation. Notably, EMT-promoting analytes such as TGF-B, IL-6, and IL-8 were consistently elevated in CAF-conditioned medium and in the medium collected from MDA-MB-231 and MCF-7 cells cultured with CAF-CM, while several analytes characteristic of epithelial states, including those secreted by MCF-7 cells, were reduced following CAF-CM treatment. This dual pattern, upregulation of EMT-inducing signals together with suppression of epithelial-associated factors, provides evidence that CAF-derived secretions remodel the tumour microenvironment in a way that promotes epithelial plasticity and drives cancer cells towards mesenchymal-like phenotypes.

In summary, this study establishes CellPhe as a framework for analysing phenotypic changes in cancer–stromal interactions directly from label-free time-lapse imaging. Through automated feature extraction, classification, and clustering, CellPhe enabled identification of breast cancer cells within co-cultures, allowing their phenotypes to be directly compared with monoculture breast cancer cells and revealing hallmarks of EMT driven by both contact and soluble CAF signals. The toolkit generalises across cell types and extends to complex multicellular systems, providing a scalable means to study tumour–microenvironment interactions. Coupling imaging-based phenotyping with immunoassays further linked phenotypes to CAF-derived EMT factors, demonstrating how CellPhe can generate mechanistic hypotheses for molecular validation.

## Acknowledgements

The authors would like to acknowledge Prof Valerie Speirs, who kindly shared the CAF cell line used in this study. The authors also gratefully acknowledge assistance from the Imaging and Cytometry Lab in the Bioscience Technology Facility at the University of York. This work was supported by the MRC (MR/X018067/1), BBSRC (BB/Y513970/1), and the Wellcome Trust (310891/Z/24/Z).

## Authors’ contributions

LW, JM, KH, PO’T, JW and WB contributed to the conception and design of the work. LW, JM, KH, PO’T, JW and WB contributed to acquisition, analysis, and interpretation of data for the work. LW, JM and WB contributed to drafting the work and revising it critically for important intellectual content. All authors approved the final version of the manuscript.

