## Supplementary Data for "Characterising cancer-stroma interactions through high-content phenotyping from microscopy time-lapses"

**Supplementary Table 1. Phenotypic features with highest separation scores for MDA-MB-231 vs. CAF**

| Feature | Separation | Category |
|---|---|---|
| Sphericity_mean | 1.657512938 | shape |
| Cooc02Pro_des | 1.424601529 | texture |
| Cooc12Pro_des | 1.367492486 | texture |
| Cooc02Pro_asc | 1.333763498 | texture |
| Cooc12Pro_asc | 1.286627117 | texture |
| Cooc01Pro_des | 1.243199116 | texture |
| dens_max | 1.240182173 | density |
| IQ7_mean | 1.227919307 | texture |
| IQ8_mean | 1.183004485 | texture |
| Sphericity_max | 1.18226362 | shape |
| Cooc01Pro_asc | 1.180900188 | texture |
| IQ6_mean | 1.157202479 | texture |
| dens_asc | 1.135952526 | density |
| Cooc02Var_des | 1.126346526 | texture |
| Cooc02Var_asc | 1.066557699 | texture |
| Cooc12Var_des | 1.05864433 | texture |
| dens_des | 1.045302062 | density |
| IQ5_mean | 1.044861835 | texture |
| Cooc01Var_des | 1.036388514 | texture |
| Sphericity_des | 1.012511903 | shape |
| Cooc12Var_asc | 1.002260845 | texture |
| Cooc01Var_asc | 0.9840442971 | texture |
| Sphericity_asc | 0.9799825529 | shape |
| dens_std | 0.978890014 | density |
| dens_mean | 0.9759710167 | density |
| Cooc12Pro_std | 0.9651540053 | texture |
| IQ9_mean | 0.9291970668 | texture |
| IQ4_mean | 0.9243405052 | texture |
| Cooc02Pro_std | 0.8690458527 | texture |
| Cooc02Pro_l1_des | 0.8564242561 | texture |
| dens_l1_des | 0.824101258 | density |
| IQ3_mean | 0.8188624457 | texture |
| Sphericity_l1_asc | 0.8174552522 | shape |
| IQ8_des | 0.8023920966 | texture |

| | | |
|---|---|---|
| Sphericity_l1_des | 0.8009012502 | shape |
| Cooc02Pro_l1_asc | 0.800222249 | texture |
| Cooc12Pro_l1_asc | 0.7991174625 | texture |
| Cooc01Pro_l1_des | 0.7933787921 | texture |
| IQ8_asc | 0.7926322239 | texture |
| Cooc12Pro_l1_des | 0.7844741763 | texture |
| IQ7_des | 0.7802284482 | texture |
| IQ7_asc | 0.7752709389 | texture |
| dens_l2_des | 0.775256691 | density |
| Cooc02Var_std | 0.773475162 | texture |
| dens_l1_asc | 0.7561428818 | density |
| Sphericity_l2_asc | 0.7560649176 | shape |
| Cooc12Var_std | 0.7479242561 | texture |
| dens_l2_asc | 0.7466950215 | density |
| Len_mean | 0.7437216246 | size |
| IQ2_mean | 0.7387565939 | texture |
| Cooc02Pro_l2_des | 0.7359047455 | texture |
| Rad_mean | 0.7356949635 | size |
| Cooc01Pro_l1_asc | 0.7314380163 | texture |
| Cooc12Pro_l2_des | 0.7132986472 | texture |
| Cooc01Var_std | 0.7115620276 | texture |
| Cooc12Pro_l2_asc | 0.7104218546 | texture |
| Cooc02Pro_l2_asc | 0.7050879021 | texture |
| IQ8_max | 0.7037840632 | texture |
| Cooc02Var_l1_des | 0.6971986691 | texture |
| Cooc01Pro_std | 0.6893768085 | texture |
| Sphericity_l2_des | 0.6841924554 | shape |
| IQ9_des | 0.6829229457 | texture |
| IQ1_mean | 0.6826996463 | texture |
| IQ9_asc | 0.675485717 | texture |
| IQ7_max | 0.6733059403 | texture |
| Sphericity_std | 0.6591469468 | shape |
| Cooc01Var_l1_des | 0.644296282 | texture |
| Cooc01Pro_l2_asc | 0.6442193327 | texture |
| Cooc01Pro_l2_des | 0.6389166404 | texture |
| IQ8_std | 0.6373671898 | texture |
| IQ9_max | 0.63358069 | texture |

| | | |
|---|---|---|
| IQ9_std | 0.6235993138 | texture |
| IQ6_des | 0.6204035167 | texture |
| IQ7_std | 0.6203502905 | texture |
| Cooc12Var_l1_des | 0.6197488419 | texture |
| IQ6_max | 0.6183955028 | texture |
| Cooc02Var_l1_asc | 0.6104744387 | texture |
| Cooc12Pro_l1_max | 0.6102768912 | texture |
| dens_l3_des | 0.6050979277 | density |
| IQ6_asc | 0.6023651585 | texture |
| Cooc12Var_l1_asc | 0.5963370049 | texture |
| Cooc01Sha_des | 0.594722656 | texture |
| IQ7_l1_asc | 0.593060447 | texture |
| IQ8_l1_des | 0.5907502981 | texture |
| IQ8_l1_asc | 0.5872742152 | texture |
| Cooc02Pro_l1_max | 0.5803103917 | texture |
| IQ7_l1_des | 0.5799311808 | texture |
| Cooc12Sha_des | 0.5791361149 | texture |
| dens_l1_max | 0.568161415 | density |
| Cooc02Var_l2_asc | 0.5650101941 | texture |
| Cooc12Var_max | 0.5607747131 | texture |
| Cooc02Var_l2_des | 0.5607584193 | texture |
| IQ5_max | 0.5607390212 | texture |
| Cooc01Var_l1_asc | 0.5584957973 | texture |
| Cooc02Sha_des | 0.5556609661 | texture |
| Rad_max | 0.5536815964 | size |
| IQ6_std | 0.5479816401 | texture |
| Cooc01Sha_asc | 0.5424414205 | texture |
| dens_l2_max | 0.5348167103 | density |
| Box_mean | 0.5346942134 | shape |
| Cooc12Sha_asc | 0.5334901534 | texture |
| Cooc12Var_l2_asc | 0.5281858097 | texture |
| Cooc12Pro_l2_max | 0.5272129876 | texture |
| Cooc01Var_l2_asc | 0.5247192838 | texture |
| IQ9_l1_asc | 0.5228976895 | texture |
| Cooc01Var_max | 0.5217157803 | texture |
| IQ4_max | 0.5151882846 | texture |
| IQ8_l2_asc | 0.5151425256 | texture |

| | | |
|---|---|---|
| Cooc01Var_I2_des | 0.5136574248 | texture |
| Cooc02Var_max | 0.5132595742 | texture |
| VfC_mean | 0.5125142751 | shape |
| Cooc02Sha_asc | 0.5097091728 | texture |
| Cooc01Pro_I1_max | 0.5091246703 | texture |
| Cooc12Var_I2_des | 0.5072390334 | texture |
| IQ8_I2_des | 0.5070131786 | texture |
| Len_max | 0.5065169283 | size |
| IQ9_I2_asc | 0.5035214538 | texture |
| dens_I3_asc | 0.502896491 | density |
| Cooc02Pro_I2_max | 0.5000328333 | texture |
| Area_mean | 0.4966879045 | size |
| IQ7_I2_des | 0.4887064991 | texture |
| IQ3_max | 0.4791342533 | texture |
| IQ9_I1_des | 0.4777961679 | texture |
| IQ6_I1_asc | 0.4753850697 | texture |
| poly1_mean | 0.473382749 | density |
| Sphericity_I3_asc | 0.469049487 | shape |
| IQ5_des | 0.4681339574 | texture |
| A2B_des | 0.4646411778 | shape |
| IQ9_I2_des | 0.4622539996 | texture |
| Cooc12Ent_asc | 0.4607788612 | texture |
| IQ5_std | 0.457650926 | texture |
| Cooc12Ent_des | 0.4574801251 | texture |
| Cooc12Sha_std | 0.4563752513 | texture |
| IQ7_I2_asc | 0.4521051741 | texture |
| IQ6_I1_des | 0.4518361167 | texture |
| IQ5_asc | 0.4517052043 | texture |
| dens_I3_max | 0.4482267235 | density |
| Cooc12Var_I1_max | 0.4474204983 | texture |
| Cooc02Ent_asc | 0.4458987843 | texture |
| IQ2_max | 0.4442808032 | texture |
| Cooc01Pro_I2_max | 0.4429299082 | texture |
| Cooc02Var_I1_max | 0.4423658333 | texture |
| Cooc02Ent_des | 0.4419955007 | texture |
| Cooc12Sha_I1_asc | 0.4418605893 | texture |
| Cooc01Sha_I1_asc | 0.4416529681 | texture |

| | | |
|---|---|---|
| Sphericity_l1_max | 0.4397433912 | shape |
| A2B_asc | 0.43627666 | shape |
| Cooc12Pro_l3_asc | 0.4337380986 | texture |
| Cooc02Pro_l3_des | 0.4326976193 | texture |
| Cooc01Sha_l1_des | 0.4325577533 | texture |
| IQ9_l3_des | 0.4307764439 | texture |
| Sphericity_l3_des | 0.4281522359 | shape |
| Cooc12Pro_l3_des | 0.4255168487 | texture |
| Cooc12Pro_max | 0.4253879143 | texture |
| Cooc02Pro_l3_asc | 0.4237057794 | texture |
| Cooc02Sha_l1_asc | 0.4196228313 | texture |
| Area_max | 0.4173894259 | size |
| Cooc02Var_l2_max | 0.4118842441 | texture |
| IQ1_max | 0.4118322164 | texture |

**Supplementary Table 2. Phenotypic features with highest separation scores for MCF-7 vs. CAF**

| Feature | Separation | Category |
|---|---|---|
| dens_max | 2.783068473 | density |
| dens_mean | 2.605707811 | density |
| Rect_mean | 2.262055985 | shape |
| Sphericity_max | 2.143691194 | shape |
| IQ4_mean | 2.076460806 | texture |
| IQ5_mean | 2.070355067 | texture |
| IQ3_mean | 2.041281338 | texture |
| IQ6_mean | 2.002791424 | texture |
| IQ2_mean | 1.977504334 | texture |
| Sphericity_mean | 1.917401225 | shape |
| IQ2_max | 1.914153608 | texture |
| IQ3_max | 1.911414488 | texture |
| dens_asc | 1.908445443 | density |
| IQ1_mean | 1.891500425 | texture |
| IQ1_max | 1.877304991 | texture |
| IQ7_mean | 1.874890443 | texture |
| IQ4_max | 1.865118125 | texture |
| IQ4_std | 1.847686199 | texture |
| IQ3_std | 1.815464491 | texture |

| | | |
|---|---|---|
| IQ5_std | 1.813801227 | texture |
| IQ5_max | 1.773590301 | texture |
| dens_des | 1.755133891 | density |
| IQ2_std | 1.754144683 | texture |
| A2B_des | 1.739012406 | shape |
| Len_max | 1.692304427 | size |
| A2B_asc | 1.691842297 | shape |
| Len_mean | 1.683630621 | size |
| IQ1_std | 1.682126253 | texture |
| IQ8_mean | 1.669010132 | texture |
| IQ6_std | 1.637967322 | texture |
| Rect_max | 1.637585506 | shape |
| Curv_asc | 1.590389495 | shape |
| Curv_des | 1.575012063 | shape |
| IQ6_max | 1.560040575 | texture |
| dens_l1_des | 1.552189804 | density |
| IQ5_des | 1.434112676 | texture |
| IQ5_asc | 1.429731325 | texture |
| IQ6_asc | 1.419449918 | texture |
| poly1_mean | 1.414875897 | density |
| IQ6_des | 1.414412052 | texture |
| Rad_max | 1.408511292 | size |
| dens_l1_asc | 1.402071934 | density |
| IQ7_std | 1.344631816 | texture |
| IQ4_asc | 1.322421266 | texture |
| IQ4_des | 1.316643163 | texture |
| dens_l2_des | 1.300710186 | density |
| dens_std | 1.29851428 | density |
| Len_std | 1.281169334 | size |
| IQ7_asc | 1.259688974 | texture |
| IQ7_des | 1.257915531 | texture |
| IQ9_mean | 1.250468705 | texture |
| IQ7_max | 1.250238693 | texture |
| Rad_mean | 1.20226273 | size |
| IQ3_asc | 1.19060965 | texture |
| A2B_max | 1.188675583 | shape |
| A2B_mean | 1.177106426 | shape |

| | | |
|---|---|---|
| poly1_max | 1.174354633 | density |
| poly1_std | 1.173043295 | density |
| Sphericity_asc | 1.170369872 | shape |
| IQ3_des | 1.167240951 | texture |
| Sphericity_des | 1.163293304 | shape |
| VfC_max | 1.152190778 | shape |
| VfC_mean | 1.121734267 | shape |
| dens_l1_max | 1.111942428 | density |
| IQ6_l1_asc | 1.104757493 | texture |
| dens_l2_asc | 1.09530444 | density |
| IQ2_asc | 1.089232607 | texture |
| IQ5_l1_asc | 1.086703 | texture |
| VfC_std | 1.078842378 | shape |
| IQ2_des | 1.058931538 | texture |
| IQ8_std | 1.058031529 | texture |
| dens_l2_max | 1.050285591 | density |
| Box_mean | 1.049770572 | shape |
| Cooc02Pro_des | 1.043334362 | texture |
| IQ5_l2_des | 1.034869745 | texture |
| IQ5_l1_des | 1.030728272 | texture |
| IQ6_l2_des | 1.027605646 | texture |
| Cooc02Pro_asc | 1.02670679 | texture |
| A2B_l1_asc | 1.021483102 | shape |
| Cooc12Pro_des | 1.015533274 | texture |
| IQ6_l1_des | 1.015468319 | texture |
| Cooc12Pro_asc | 1.000101732 | texture |
| IQ1_asc | 0.994992057 | texture |
| IQ4_l2_des | 0.988154789 | texture |
| Sphericity_l1_asc | 0.980451258 | shape |
| IQ8_des | 0.98040646 | texture |
| A2B_l1_des | 0.979748131 | shape |
| IQ4_l1_des | 0.979508987 | texture |
| IQ4_l1_asc | 0.979359913 | texture |
| IQ5_l2_asc | 0.977106575 | texture |
| IQ1_des | 0.969560524 | texture |
| IQ8_asc | 0.96823394 | texture |
| IQ7_l1_asc | 0.967571467 | texture |

| | | |
|---|---|---|
| IQ6_l2_asc | 0.964004298 | texture |
| IQ5_l3_des | 0.961682079 | texture |
| Curv_max | 0.960160176 | shape |
| IQ8_max | 0.95961313 | texture |
| Cooc01Pro_des | 0.952756549 | texture |
| Cooc01Pro_asc | 0.950096297 | texture |
| IQ7_l2_des | 0.940118556 | texture |
| IQ4_l3_des | 0.939295168 | texture |
| IQ7_l1_des | 0.937662701 | texture |
| Sphericity_l1_des | 0.936617676 | shape |
| IQ5_l3_asc | 0.929310903 | texture |
| IQ6_l3_des | 0.926929487 | texture |
| IQ3_l2_des | 0.9255833 | texture |
| IQ6_l3_asc | 0.916486355 | texture |
| IQ3_l3_des | 0.912620748 | texture |
| Rad_std | 0.904937075 | size |
| IQ4_l3_asc | 0.904030673 | texture |
| Cooc12Var_des | 0.897835163 | texture |
| IQ7_l2_asc | 0.891049745 | texture |
| Cooc12Var_asc | 0.890713154 | texture |
| IQ3_l1_des | 0.889475346 | texture |
| Cooc02Var_des | 0.878089608 | texture |
| IQ4_l2_asc | 0.87699883 | texture |
| IQ2_l2_des | 0.876968341 | texture |
| VfC_asc | 0.875104236 | shape |
| Cooc02Var_asc | 0.870509774 | texture |
| IQ2_l3_des | 0.868747175 | texture |
| Curv_std | 0.867074964 | shape |
| IQ3_l1_asc | 0.865175673 | texture |
| Cooc02Sva_max | 0.855008174 | texture |
| dens_l3_des | 0.847691673 | density |
| VfC_des | 0.845177067 | shape |
| IQ9_std | 0.842444235 | texture |
| Curv_l1_asc | 0.842206783 | shape |
| Box_max | 0.840868936 | shape |
| Cooc01Var_des | 0.840233187 | texture |
| IQ3_l3_asc | 0.840036118 | texture |

| | | |
|---|---|---|
| Cooc01Sva_max | 0.838489487 | texture |
| IQ7_I3_asc | 0.83847035 | texture |
| Cooc01Var_asc | 0.835381475 | texture |
| IQ1_I3_des | 0.825591507 | texture |
| IQ1_I2_des | 0.825267562 | texture |
| Box_std | 0.821061466 | shape |
| IQ2_I1_des | 0.809914137 | texture |
| IQ2_I1_asc | 0.793478602 | texture |
| IQ7_I3_des | 0.791667486 | texture |
| IQ3_I2_asc | 0.78844778 | texture |
| FOmean_skew | 0.783511758 | texture |
| VfC_I2_des | 0.778186791 | shape |
| Sphericity_I2_asc | 0.774156039 | shape |
| Rad_des | 0.772884444 | size |
| Curv_I1_des | 0.772213913 | shape |
| IQ8_I2_asc | 0.768394367 | texture |
| Cooc02Pro_I1_des | 0.765212594 | texture |
| Cooc12Var_max | 0.758482713 | texture |
| IQ2_I3_asc | 0.758455881 | texture |
| Cooc12Pro_I1_asc | 0.757847782 | texture |
| Sphericity_I2_des | 0.757596936 | shape |
| Cooc02Pro_I1_asc | 0.75719595 | texture |
| VfC_I1_des | 0.755767063 | shape |

**Supplementary Table 3. Directionality of significant phenotypic features for MDA-MB-231 cells cultured with increasing density of CAFs.**

| Correlation Direction | Number of Features | Feature Names |
|---|---|---|
| Positively correlated (ρ ≥ 0.20) | 22 | dens_I1_des, dens_des, dens_I3_des, A2B_des, Sphericity_des, IQ4_mean, IQ3_mean, Len_mean, IQ2_mean, IQ5_mean, IQ1_mean, Cooc01Var_des, Cooc12Var_des, IQ6_mean, Cooc12Pro_des, Cooc02Var_des, IQ7_mean, IQ8_mean, IQ7_asc, VfC_mean, IQ6_asc, IQ6_std |
| Negatively correlated (ρ ≤ -0.20) | 30 | dens_max, dens_mean, dens_std, dens_asc, dens_I2_asc, dens_I2_max, dens_I3_asc, dens_I3_max, Sphericity_I1_max, A2B_asc, Cooc12Pro_I1_max, Cooc02Pro_I1_max, Cooc12Pro_I2_max, Sphericity_max, Sphericity_asc, Cooc01Pro_I1_max, Cooc02Pro_I2_max, |

| | | |
|---|---|---|
| | | Sphericity_mean, Cooc01Pro_I2_max, Cooc12Var_I1_max, Cooc12Pro_asc, Sphericity_I1_asc, Cooc01Var_asc, Cooc02Var_I2_max, Cooc12Var_asc, Cooc02Var_asc, Cooc02Var_I1_max, Cooc02Ent_asc, IQ8_I2_des, IQ6_I1_des |

**Supplementary Table 4. Directionality of significant phenotypic features for MCF-7 cells cultured with increasing density of CAFs.**

| Correlation Direction | Number of Features | Feature Names |
|---|---|---|
| Positively correlated (ρ ≥ 0.2) | 13 | Cooc01Var_des, Cooc02Var_des, Cooc12Var_des, dens_des, Cooc02Pro_des, Cooc12Pro_des, dens_I1_des, Cooc01Pro_des, dens_I2_des, Cooc02Pro_I1_des, dens_I3_des, Sphericity_I2_des, Sphericity_I1_des |
| Negatively correlated (ρ ≤ -0.0) | 20 | Cooc01Var_asc, Cooc02Var_asc, Cooc12Var_asc, dens_max, dens_std, dens_asc, Cooc02Pro_asc, Cooc12Pro_asc, dens_I2_asc, Cooc01Pro_asc, dens_mean, dens_I1_asc, dens_I2_max, Cooc12Pro_I1_asc, Cooc02Pro_I1_asc, dens_I1_max, Sphericity_mean, Sphericity_asc, Sphericity_I2_asc, Sphericity_I1_asc |

**Supplementary Table 5. Significant phenotypic changes (p < 0.01):**

| Cell Line | Number of Significant Features | Feature Names |
|---|---|---|
| MDA-MB-231 | 787 | Volume_mean, Volume_des, Volume_I1_asc, Sphericity_mean, Sphericity_std, Sphericity_asc, Sphericity_des, Sphericity_max, Sphericity_I1_asc, Sphericity_I1_des, Sphericity_I1_max, Sphericity_I2_asc, Sphericity_I2_des, Sphericity_I2_max, Sphericity_I3_asc, Sphericity_I3_des, Sphericity_I3_max, Dis_asc, Dis_I1_des, Dis_I2_des, Trac_asc, Trac_I1_des, Trac_I2_des, Trac_I3_des, D2T_asc, D2T_des, D2T_I1_des, Vel_mean, Vel_std, Vel_asc, Vel_des, Vel_I1_des, Rad_mean, Rad_std, Rad_skew, Rad_asc, Rad_des, Rad_max, Rad_I1_asc, Rad_I1_des, Rad_I1_max, Rad_I2_asc, Rad_I2_des, Rad_I3_asc, Rad_I3_des, VfC_mean, VfC_std, VfC_skew, VfC_asc, VfC_des, VfC_max, VfC_I1_asc, VfC_I1_des, VfC_I1_max, VfC_I2_asc, VfC_I2_des, VfC_I3_asc, VfC_I3_des, VfC_I3_max, Curv_mean, Curv_std, Curv_asc, Curv_des, Curv_max, |

| | | |
|---|---|---|
| | | Curv_l1_asc, Curv_l1_des, Curv_l1_max, Curv_l2_asc, Curv_l2_des, Curv_l2_max, Curv_l3_max, Len_mean, Len_std, Len_skew, Len_asc, Len_des, Len_max, Len_l1_asc, Len_l1_des, Len_l2_asc, Len_l2_des, Len_l3_asc, Len_l3_des, Wid_std, Wid_skew, Wid_asc, Wid_des, Wid_max, Wid_l1_asc, Wid_l1_max, Wid_l2_des, Wid_l3_asc, Wid_l3_des, Wid_l3_max, Area_mean, Area_asc, Area_des, Area_max, Area_l1_asc, Area_l3_asc, A2B_mean, A2B_skew, A2B_asc, A2B_des, A2B_max, A2B_l1_asc, A2B_l1_des, A2B_l1_max, A2B_l2_asc, A2B_l2_des, A2B_l3_asc, A2B_l3_max, Box_mean, Box_std, Box_skew, Box_asc, Box_des, Box_max, Box_l1_asc, Box_l1_des, Box_l1_max, Box_l2_asc, Box_l2_des, Box_l3_asc, Box_l3_des, Rect_mean, Rect_std, Rect_skew, Rect_max, Rect_l2_max, poly1_mean, poly1_std, poly1_skew, poly1_asc, poly1_des, poly1_max, poly1_l1_asc, poly1_l1_des, poly1_l1_max, poly1_l2_asc, poly1_l2_des, poly2_mean, poly2_skew, poly2_max, poly3_mean, poly3_std, poly3_asc, poly3_des, poly3_max, poly3_l1_asc, poly3_l1_des, poly3_l2_asc, poly3_l3_asc, poly4_mean, poly4_std, poly4_asc, poly4_des, poly4_max, poly4_l1_asc, poly4_l1_des, poly4_l1_max, poly4_l2_asc, poly4_l2_des, poly4_l2_max, poly4_l3_asc, poly4_l3_des, FOmean_mean, FOmean_std, FOmean_asc, FOmean_des, FOmean_max, FOmean_l1_asc, FOmean_l1_des, FOmean_l1_max, FOmean_l2_asc, FOmean_l2_des, FOmean_l2_max, FOmean_l3_asc, FOmean_l3_des, FOmean_l3_max, FOsd_mean, FOsd_std, FOsd_asc, FOsd_des, FOsd_max, FOsd_l1_asc, FOsd_l1_des, FOsd_l1_max, FOsd_l2_asc, FOsd_l2_des, FOsd_l2_max, FOsd_l3_asc, FOsd_l3_des, FOsd_l3_max, FOskew_mean, FOskew_skew, Cooc01ASM_mean, Cooc01ASM_std, Cooc01ASM_skew, Cooc01ASM_asc, Cooc01ASM_des, Cooc01ASM_max, Cooc01ASM_l1_asc, Cooc01ASM_l1_des, Cooc01ASM_l1_max, Cooc01ASM_l2_asc, Cooc01ASM_l2_des, Cooc01ASM_l2_max, Cooc01ASM_l3_asc, Cooc01ASM_l3_des, Cooc01ASM_l3_max, Cooc01Con_mean, Cooc01Con_std, Cooc01Con_asc, Cooc01Con_des, Cooc01Con_max, Cooc01Con_l1_asc, Cooc01Con_l1_des, Cooc01Con_l1_max, Cooc01Con_l2_asc, Cooc01Con_l2_des, Cooc01Con_l2_max, Cooc01Con_l3_asc, Cooc01Con_l3_des, Cooc01Con_l3_max, Cooc01IDM_std, Cooc01IDM_asc, Cooc01IDM_des, Cooc01IDM_max, |

| | | |
|---|---|---|
| | | Cooc01IDM_l1_asc, Cooc01IDM_l1_des, Cooc01IDM_l1_max, Cooc01IDM_l2_asc, Cooc01IDM_l2_des, Cooc01IDM_l2_max, Cooc01IDM_l3_asc, Cooc01IDM_l3_des, Cooc01IDM_l3_max, Cooc01Ent_mean, Cooc01Ent_std, Cooc01Ent_skew, Cooc01Ent_asc, Cooc01Ent_des, Cooc01Ent_l1_asc, Cooc01Ent_l1_des, Cooc01Ent_l1_max, Cooc01Ent_l2_asc, Cooc01Ent_l2_des, Cooc01Ent_l2_max, Cooc01Ent_l3_asc, Cooc01Ent_l3_des, Cooc01Ent_l3_max, Cooc01Cor_mean, Cooc01Cor_std, Cooc01Cor_asc, Cooc01Cor_des, Cooc01Cor_max, Cooc01Cor_l1_asc, Cooc01Cor_l1_des, Cooc01Cor_l1_max, Cooc01Cor_l2_asc, Cooc01Cor_l2_des, Cooc01Cor_l2_max, Cooc01Cor_l3_asc, Cooc01Cor_l3_des, Cooc01Cor_l3_max, Cooc01Var_mean, Cooc01Var_std, Cooc01Var_max, Cooc01Var_l3_des, Cooc01Sav_mean, Cooc01Sav_std, Cooc01Sav_skew, Cooc01Sav_asc, Cooc01Sav_des, Cooc01Sav_max, Cooc01Sav_l1_des, Cooc01Sav_l1_max, Cooc01Sav_l2_asc, Cooc01Sav_l3_asc, Cooc01Sen_mean, Cooc01Sen_std, Cooc01Sen_asc, Cooc01Sen_des, Cooc01Sen_max, Cooc01Sen_l1_asc, Cooc01Sen_l1_des, Cooc01Sen_l1_max, Cooc01Sen_l2_asc, Cooc01Sen_l2_des, Cooc01Sen_l2_max, Cooc01Sen_l3_asc, Cooc01Sen_l3_des, Cooc01Sen_l3_max, Cooc01Den_std, Cooc01Den_skew, Cooc01Den_asc, Cooc01Den_des, Cooc01Den_max, Cooc01Den_l1_asc, Cooc01Den_l1_des, Cooc01Den_l1_max, Cooc01Den_l2_asc, Cooc01Den_l2_des, Cooc01Den_l2_max, Cooc01Den_l3_asc, Cooc01Den_l3_des, Cooc01Den_l3_max, Cooc01Dva_mean, Cooc01Dva_std, Cooc01Dva_asc, Cooc01Dva_des, Cooc01Dva_max, Cooc01Dva_l1_asc, Cooc01Dva_l1_des, Cooc01Dva_l1_max, Cooc01Dva_l2_asc, Cooc01Dva_l2_des, Cooc01Dva_l3_asc, Cooc01Dva_l3_des, Cooc01Dva_l3_max, Cooc01Sva_mean, Cooc01Sva_skew, Cooc01Sva_max, Cooc01f13_mean, Cooc01f13_std, Cooc01f13_asc, Cooc01f13_des, Cooc01f13_max, Cooc01f13_l1_asc, Cooc01f13_l1_des, Cooc01f13_l1_max, Cooc01f13_l2_asc, Cooc01f13_l2_des, Cooc01f13_l2_max, Cooc01f13_l3_asc, Cooc01f13_l3_des, Cooc01f13_l3_max, Cooc01Sha_skew, Cooc01Sha_asc, Cooc01Sha_des, Cooc01Sha_l1_des, Cooc01Pro_mean, |

| | | |
|---|---|---|
| | | Cooc01Pro_max, Cooc12ASM_mean, Cooc12ASM_std, Cooc12ASM_skew, Cooc12ASM_asc, Cooc12ASM_des, Cooc12ASM_max, Cooc12ASM_l1_asc, Cooc12ASM_l1_des, Cooc12ASM_l1_max, Cooc12ASM_l2_des, Cooc12ASM_l3_asc, Cooc12ASM_l3_des, Cooc12ASM_l3_max, Cooc12Con_std, Cooc12Con_asc, Cooc12Con_des, Cooc12Con_max, Cooc12Con_l1_asc, Cooc12Con_l1_des, Cooc12Con_l1_max, Cooc12Con_l2_asc, Cooc12Con_l2_des, Cooc12Con_l2_max, Cooc12Con_l3_asc, Cooc12Con_l3_des, Cooc12Con_l3_max, Cooc12IDM_std, Cooc12IDM_asc, Cooc12IDM_des, Cooc12IDM_max, Cooc12IDM_l1_asc, Cooc12IDM_l1_des, Cooc12IDM_l1_max, Cooc12IDM_l2_asc, Cooc12IDM_l2_des, Cooc12IDM_l2_max, Cooc12IDM_l3_asc, Cooc12IDM_l3_des, Cooc12IDM_l3_max, Cooc12Ent_mean, Cooc12Ent_std, Cooc12Ent_skew, Cooc12Ent_asc, Cooc12Ent_des, Cooc12Ent_l1_asc, Cooc12Ent_l1_des, Cooc12Ent_l1_max, Cooc12Ent_l2_asc, Cooc12Ent_l2_des, Cooc12Ent_l2_max, Cooc12Ent_l3_asc, Cooc12Ent_l3_des, Cooc12Ent_l3_max, Cooc12Cor_mean, Cooc12Cor_std, Cooc12Cor_skew, Cooc12Cor_asc, Cooc12Cor_des, Cooc12Cor_max, Cooc12Cor_l1_asc, Cooc12Cor_l1_des, Cooc12Cor_l1_max, Cooc12Cor_l2_asc, Cooc12Cor_l2_des, Cooc12Cor_l2_max, Cooc12Cor_l3_asc, Cooc12Cor_l3_des, Cooc12Cor_l3_max, Cooc12Var_mean, Cooc12Var_std, Cooc12Var_max, Cooc12Var_l3_des, Cooc12Sav_mean, Cooc12Sav_std, Cooc12Sav_skew, Cooc12Sav_asc, Cooc12Sav_des, Cooc12Sav_max, Cooc12Sav_l1_des, Cooc12Sav_l1_max, Cooc12Sav_l2_asc, Cooc12Sav_l3_asc, Cooc12Sav_l3_des, Cooc12Sav_l3_max, Cooc12Sen_mean, Cooc12Sen_std, Cooc12Sen_asc, Cooc12Sen_des, Cooc12Sen_max, Cooc12Sen_l1_asc, Cooc12Sen_l1_des, Cooc12Sen_l1_max, Cooc12Sen_l2_asc, Cooc12Sen_l2_des, Cooc12Sen_l2_max, Cooc12Sen_l3_asc, Cooc12Sen_l3_des, Cooc12Sen_l3_max, Cooc12Den_std, Cooc12Den_skew, Cooc12Den_asc, Cooc12Den_des, Cooc12Den_max, Cooc12Den_l1_asc, Cooc12Den_l1_des, Cooc12Den_l1_max, Cooc12Den_l2_asc, Cooc12Den_l2_des, Cooc12Den_l2_max, Cooc12Den_l3_asc, Cooc12Den_l3_des, Cooc12Den_l3_max, Cooc12Dva_mean, |

| | | |
|---|---|---|
| | | Cooc12Dva_std, Cooc12Dva_asc, Cooc12Dva_des, Cooc12Dva_max, Cooc12Dva_l1_asc, Cooc12Dva_l1_des, Cooc12Dva_l1_max, Cooc12Dva_l2_asc, Cooc12Dva_l2_des, Cooc12Dva_l2_max, Cooc12Dva_l3_asc, Cooc12Dva_l3_des, Cooc12Dva_l3_max, Cooc12Sva_mean, Cooc12Sva_std, Cooc12Sva_skew, Cooc12Sva_max, Cooc12Sva_l1_des, Cooc12Sva_l3_asc, Cooc12f13_mean, Cooc12f13_std, Cooc12f13_skew, Cooc12f13_asc, Cooc12f13_des, Cooc12f13_max, Cooc12f13_l1_asc, Cooc12f13_l1_des, Cooc12f13_l1_max, Cooc12f13_l2_asc, Cooc12f13_l2_des, Cooc12f13_l2_max, Cooc12f13_l3_asc, Cooc12f13_l3_des, Cooc12f13_l3_max, Cooc12Sha_skew, Cooc12Sha_asc, Cooc12Sha_des, Cooc12Sha_l1_des, Cooc12Pro_mean, Cooc12Pro_max, Cooc02ASM_mean, Cooc02ASM_std, Cooc02ASM_skew, Cooc02ASM_asc, Cooc02ASM_des, Cooc02ASM_max, Cooc02ASM_l1_asc, Cooc02ASM_l1_des, Cooc02ASM_l1_max, Cooc02ASM_l2_des, Cooc02ASM_l3_asc, Cooc02ASM_l3_des, Cooc02ASM_l3_max, Cooc02Con_mean, Cooc02Con_std, Cooc02Con_skew, Cooc02Con_asc, Cooc02Con_des, Cooc02Con_max, Cooc02Con_l1_asc, Cooc02Con_l1_des, Cooc02Con_l1_max, Cooc02Con_l2_asc, Cooc02Con_l2_des, Cooc02Con_l2_max, Cooc02Con_l3_asc, Cooc02Con_l3_des, Cooc02Con_l3_max, Cooc02IDM_std, Cooc02IDM_asc, Cooc02IDM_des, Cooc02IDM_max, Cooc02IDM_l1_asc, Cooc02IDM_l1_des, Cooc02IDM_l1_max, Cooc02IDM_l2_asc, Cooc02IDM_l2_des, Cooc02IDM_l2_max, Cooc02IDM_l3_asc, Cooc02IDM_l3_des, Cooc02IDM_l3_max, Cooc02Ent_mean, Cooc02Ent_std, Cooc02Ent_skew, Cooc02Ent_asc, Cooc02Ent_des, Cooc02Ent_l1_asc, Cooc02Ent_l1_des, Cooc02Ent_l1_max, Cooc02Ent_l2_asc, Cooc02Ent_l2_des, Cooc02Ent_l2_max, Cooc02Ent_l3_asc, Cooc02Ent_l3_des, Cooc02Ent_l3_max, Cooc02Cor_mean, Cooc02Cor_std, Cooc02Cor_skew, Cooc02Cor_asc, Cooc02Cor_des, Cooc02Cor_max, Cooc02Cor_l1_asc, Cooc02Cor_l1_des, Cooc02Cor_l1_max, Cooc02Cor_l2_asc, Cooc02Cor_l2_des, Cooc02Cor_l2_max, Cooc02Cor_l3_asc, Cooc02Cor_l3_des, Cooc02Cor_l3_max, Cooc02Var_mean, Cooc02Var_std, Cooc02Var_max, Cooc02Var_l3_des, |

| | | |
|---|---|---|
| | | Cooc02Sav_mean, Cooc02Sav_std, Cooc02Sav_skew, Cooc02Sav_asc, Cooc02Sav_des, Cooc02Sav_max, Cooc02Sav_l1_des, Cooc02Sav_l1_max, Cooc02Sav_l2_asc, Cooc02Sav_l3_asc, Cooc02Sav_l3_des, Cooc02Sen_mean, Cooc02Sen_std, Cooc02Sen_asc, Cooc02Sen_des, Cooc02Sen_max, Cooc02Sen_l1_asc, Cooc02Sen_l1_des, Cooc02Sen_l1_max, Cooc02Sen_l2_asc, Cooc02Sen_l2_des, Cooc02Sen_l2_max, Cooc02Sen_l3_asc, Cooc02Sen_l3_des, Cooc02Sen_l3_max, Cooc02Den_std, Cooc02Den_asc, Cooc02Den_des, Cooc02Den_max, Cooc02Den_l1_asc, Cooc02Den_l1_des, Cooc02Den_l1_max, Cooc02Den_l2_asc, Cooc02Den_l2_des, Cooc02Den_l2_max, Cooc02Den_l3_asc, Cooc02Den_l3_des, Cooc02Den_l3_max, Cooc02Dva_mean, Cooc02Dva_std, Cooc02Dva_skew, Cooc02Dva_asc, Cooc02Dva_des, Cooc02Dva_max, Cooc02Dva_l1_asc, Cooc02Dva_l1_des, Cooc02Dva_l1_max, Cooc02Dva_l2_asc, Cooc02Dva_l2_des, Cooc02Dva_l2_max, Cooc02Dva_l3_asc, Cooc02Dva_l3_des, Cooc02Dva_l3_max, Cooc02Sva_mean, Cooc02Sva_std, Cooc02Sva_skew, Cooc02Sva_max, Cooc02f13_mean, Cooc02f13_std, Cooc02f13_skew, Cooc02f13_asc, Cooc02f13_des, Cooc02f13_max, Cooc02f13_l1_asc, Cooc02f13_l1_des, Cooc02f13_l1_max, Cooc02f13_l2_asc, Cooc02f13_l2_des, Cooc02f13_l2_max, Cooc02f13_l3_asc, Cooc02f13_l3_des, Cooc02f13_l3_max, Cooc02Sha_skew, Cooc02Sha_asc, Cooc02Sha_des, Cooc02Sha_l1_des, Cooc02Pro_mean, Cooc02Pro_max, IQ1_mean, IQ1_std, IQ1_skew, IQ1_asc, IQ1_des, IQ1_max, IQ1_l1_asc, IQ1_l1_des, IQ1_l1_max, IQ1_l2_asc, IQ1_l2_des, IQ1_l3_asc, IQ1_l3_des, IQ2_mean, IQ2_std, IQ2_skew, IQ2_asc, IQ2_des, IQ2_max, IQ2_l1_asc, IQ2_l1_des, IQ2_l1_max, IQ2_l2_asc, IQ2_l2_des, IQ2_l3_asc, IQ2_l3_des, IQ3_mean, IQ3_std, IQ3_skew, IQ3_asc, IQ3_des, IQ3_max, IQ3_l1_asc, IQ3_l1_des, IQ3_l1_max, IQ3_l2_asc, IQ3_l2_des, IQ3_l3_asc, IQ3_l3_des, IQ3_l3_max, IQ4_mean, IQ4_std, IQ4_skew, IQ4_asc, IQ4_des, IQ4_max, IQ4_l1_asc, IQ4_l1_des, IQ4_l1_max, IQ4_l2_asc, IQ4_l2_des, IQ4_l3_asc, IQ4_l3_des, IQ4_l3_max, IQ5_mean, IQ5_std, IQ5_skew, IQ5_asc, IQ5_des, IQ5_max, IQ5_l1_asc, IQ5_l1_des, IQ5_l1_max, IQ5_l2_asc, IQ5_l2_des, IQ5_l2_max, IQ5_l3_asc, IQ5_l3_des, IQ5_l3_max, IQ6_mean, IQ6_std, IQ6_skew, IQ6_asc, |

| | | |
|---|---|---|
| | | IQ6_des, IQ6_max, IQ6_l1_asc, IQ6_l1_des, IQ6_l1_max, IQ6_l2_asc, IQ6_l2_des, IQ6_l2_max, IQ6_l3_asc, IQ6_l3_des, IQ6_l3_max, IQ7_mean, IQ7_std, IQ7_asc, IQ7_des, IQ7_max, IQ7_l1_asc, IQ7_l1_des, IQ7_l1_max, IQ7_l2_asc, IQ7_l2_des, IQ7_l2_max, IQ7_l3_asc, IQ7_l3_des, IQ7_l3_max, IQ8_mean, IQ8_std, IQ8_skew, IQ8_asc, IQ8_des, IQ8_max, IQ8_l1_asc, IQ8_l1_des, IQ8_l1_max, IQ8_l2_asc, IQ8_l2_des, IQ8_l2_max, IQ8_l3_asc, IQ8_l3_des, IQ8_l3_max, IQ9_mean, IQ9_std, IQ9_asc, IQ9_des, IQ9_max, IQ9_l1_asc, IQ9_l1_des, IQ9_l1_max, IQ9_l2_asc, IQ9_l2_des, IQ9_l2_max, IQ9_l3_asc, IQ9_l3_des, IQ9_l3_max, dens_mean, dens_std, dens_skew, dens_asc, dens_des, dens_max, dens_l1_asc, dens_l1_des, dens_l2_des, dens_l3_des, trajArea |
| MCF-7 | 566 | Volume_mean, Volume_std, Volume_skew, Volume_des, Volume_max, Sphericity_mean, Sphericity_std, Sphericity_skew, Sphericity_asc, Sphericity_des, Sphericity_max, Sphericity_l1_asc, Sphericity_l1_des, Sphericity_l1_max, Sphericity_l2_asc, Sphericity_l2_des, Sphericity_l3_asc, Sphericity_l3_des, Dis_mean, Dis_std, Dis_skew, Dis_max, Dis_l2_max, Dis_l3_des, Dis_l3_max, Trac_mean, Trac_std, Trac_max, D2T_mean, D2T_skew, D2T_asc, D2T_des, D2T_l1_des, Rad_std, Rad_max, Rad_l2_max, VfC_mean, VfC_std, VfC_asc, VfC_des, VfC_max, VfC_l1_asc, VfC_l1_max, VfC_l2_asc, VfC_l2_max, VfC_l3_asc, VfC_l3_des, VfC_l3_max, Curv_mean, Curv_std, Curv_skew, Curv_asc, Curv_des, Curv_max, Curv_l1_des, Curv_l1_max, Curv_l3_max, Len_mean, Len_std, Len_max, Len_l2_max, Wid_mean, Wid_std, Wid_skew, Wid_max, Wid_l2_des, Area_std, A2B_mean, A2B_std, A2B_skew, A2B_asc, A2B_des, A2B_l1_asc, A2B_l1_des, A2B_l1_max, Box_skew, Box_asc, Box_des, Box_l1_asc, Box_l1_des, Rect_skew, poly1_mean, poly1_asc, poly1_des, poly1_l1_des, poly1_l2_asc, poly2_mean, poly2_std, poly2_skew, poly2_asc, poly2_des, poly2_max, poly2_l1_asc, poly2_l1_des, poly2_l1_max, poly2_l2_asc, poly2_l2_des, poly2_l3_asc, poly3_mean, poly3_max, poly3_l3_max, poly4_mean, poly4_std, poly4_asc, poly4_des, poly4_max, poly4_l1_asc, poly4_l1_des, poly4_l1_max, poly4_l2_asc, poly4_l2_des, poly4_l2_max, poly4_l3_asc, poly4_l3_des, poly4_l3_max, FOmean_std, FOmean_skew, FOmean_asc, FOmean_des, FOmean_max, FOmean_l1_asc, FOmean_l1_des, FOmean_l2_asc, FOmean_l2_des, FOmean_l2_max, FOmean_l3_asc, |

| | | |
|---|---|---|
| | | FOmean_l3_des, FOmean_l3_max, FOsd_mean, FOsd_std, FOsd_asc, FOsd_des, FOsd_max, FOsd_l1_asc, FOsd_l1_des, FOsd_l2_asc, FOsd_l2_des, FOsd_l2_max, FOsd_l3_asc, FOsd_l3_des, FOsd_l3_max, FOskew_mean, FOskew_std, FOskew_skew, FOskew_asc, FOskew_des, FOskew_max, FOskew_l1_asc, FOskew_l1_max, FOskew_l2_asc, FOskew_l3_asc, Cooc01ASM_mean, Cooc01ASM_skew, Cooc01ASM_max, Cooc01Con_mean, Cooc01Con_std, Cooc01Con_asc, Cooc01Con_des, Cooc01Con_max, Cooc01Con_l1_asc, Cooc01Con_l1_des, Cooc01Con_l2_asc, Cooc01Con_l2_des, Cooc01Con_l2_max, Cooc01Con_l3_asc, Cooc01Con_l3_des, Cooc01IDM_mean, Cooc01IDM_std, Cooc01IDM_asc, Cooc01IDM_des, Cooc01IDM_l1_asc, Cooc01IDM_l1_des, Cooc01IDM_l1_max, Cooc01IDM_l2_asc, Cooc01IDM_l2_des, Cooc01IDM_l2_max, Cooc01IDM_l3_asc, Cooc01IDM_l3_des, Cooc01IDM_l3_max, Cooc01Ent_skew, Cooc01Cor_mean, Cooc01Cor_std, Cooc01Cor_asc, Cooc01Cor_des, Cooc01Cor_max, Cooc01Cor_l1_asc, Cooc01Cor_l1_des, Cooc01Cor_l1_max, Cooc01Cor_l2_asc, Cooc01Cor_l2_des, Cooc01Cor_l2_max, Cooc01Cor_l3_asc, Cooc01Cor_l3_des, Cooc01Cor_l3_max, Cooc01Var_mean, Cooc01Var_skew, Cooc01Var_asc, Cooc01Var_des, Cooc01Var_max, Cooc01Var_l1_des, Cooc01Var_l1_max, Cooc01Var_l2_asc, Cooc01Var_l2_max, Cooc01Var_l3_max, Cooc01Sav_std, Cooc01Sav_skew, Cooc01Sav_asc, Cooc01Sav_max, Cooc01Sen_skew, Cooc01Den_mean, Cooc01Den_skew, Cooc01Den_asc, Cooc01Den_des, Cooc01Den_max, Cooc01Den_l1_asc, Cooc01Den_l1_des, Cooc01Den_l2_asc, Cooc01Den_l2_des, Cooc01Den_l3_des, Cooc01Dva_mean, Cooc01Dva_std, Cooc01Dva_asc, Cooc01Dva_des, Cooc01Dva_l1_asc, Cooc01Dva_l1_des, Cooc01Dva_l2_asc, Cooc01Dva_l2_des, Cooc01Dva_l3_asc, Cooc01Dva_l3_des, Cooc01Sva_mean, Cooc01Sva_std, Cooc01Sva_max, Cooc01Sva_l2_max, Cooc01f13_mean, Cooc01f13_asc, Cooc01f13_des, Cooc01f13_max, Cooc01f13_l1_asc, Cooc01f13_l1_max, Cooc01Sha_mean, Cooc01Sha_std, Cooc01Sha_asc, Cooc01Sha_des, Cooc01Sha_max, Cooc01Sha_l1_asc, Cooc01Sha_l1_des, Cooc01Sha_l1_max, Cooc01Sha_l2_asc, Cooc01Sha_l2_des, Cooc01Sha_l2_max, |

| | | |
|---|---|---|
| | | Cooc01Sha_l3_asc, Cooc01Sha_l3_des, Cooc01Sha_l3_max, Cooc01Pro_mean, Cooc01Pro_std, Cooc01Pro_skew, Cooc01Pro_asc, Cooc01Pro_des, Cooc01Pro_max, Cooc01Pro_l1_asc, Cooc01Pro_l1_des, Cooc01Pro_l1_max, Cooc01Pro_l2_asc, Cooc01Pro_l2_des, Cooc01Pro_l2_max, Cooc01Pro_l3_asc, Cooc01Pro_l3_des, Cooc01Pro_l3_max, Cooc12ASM_mean, Cooc12ASM_skew, Cooc12ASM_max, Cooc12Con_std, Cooc12Con_skew, Cooc12Con_asc, Cooc12Con_des, Cooc12Con_max, Cooc12Con_l1_asc, Cooc12Con_l1_des, Cooc12Con_l2_asc, Cooc12Con_l2_des, Cooc12Con_l2_max, Cooc12Con_l3_asc, Cooc12Con_l3_des, Cooc12IDM_mean, Cooc12IDM_std, Cooc12IDM_asc, Cooc12IDM_des, Cooc12IDM_l1_asc, Cooc12IDM_l1_des, Cooc12IDM_l1_max, Cooc12IDM_l2_asc, Cooc12IDM_l2_des, Cooc12IDM_l2_max, Cooc12IDM_l3_asc, Cooc12IDM_l3_des, Cooc12IDM_l3_max, Cooc12Ent_skew, Cooc12Cor_mean, Cooc12Cor_std, Cooc12Cor_asc, Cooc12Cor_des, Cooc12Cor_max, Cooc12Cor_l1_asc, Cooc12Cor_l1_des, Cooc12Cor_l2_asc, Cooc12Cor_l2_des, Cooc12Cor_l3_asc, Cooc12Cor_l3_des, Cooc12Cor_l3_max, Cooc12Var_mean, Cooc12Var_skew, Cooc12Var_max, Cooc12Var_l1_des, Cooc12Var_l1_max, Cooc12Var_l2_max, Cooc12Sav_skew, Cooc12Sav_asc, Cooc12Sav_des, Cooc12Sav_max, Cooc12Sav_l1_asc, Cooc12Sav_l1_des, Cooc12Sen_skew, Cooc12Den_asc, Cooc12Den_des, Cooc12Den_l1_asc, Cooc12Den_l1_des, Cooc12Den_l2_asc, Cooc12Den_l2_des, Cooc12Den_l3_des, Cooc12Dva_skew, Cooc12Dva_asc, Cooc12Dva_des, Cooc12Dva_l1_asc, Cooc12Dva_l1_des, Cooc12Dva_l2_asc, Cooc12Dva_l2_des, Cooc12Dva_l2_max, Cooc12Dva_l3_des, Cooc12Sva_std, Cooc12Sva_skew, Cooc12Sva_asc, Cooc12Sva_max, Cooc12f13_skew, Cooc12f13_max, Cooc12Sha_mean, Cooc12Sha_std, Cooc12Sha_skew, Cooc12Sha_asc, Cooc12Sha_des, Cooc12Sha_max, Cooc12Sha_l1_asc, Cooc12Sha_l1_des, Cooc12Sha_l1_max, Cooc12Sha_l2_asc, Cooc12Sha_l2_des, Cooc12Sha_l2_max, Cooc12Sha_l3_des, Cooc12Sha_l3_max, Cooc12Pro_mean, Cooc12Pro_std, Cooc12Pro_skew, Cooc12Pro_asc, Cooc12Pro_des, Cooc12Pro_max, |

| | | |
|---|---|---|
| | | Cooc12Pro_l1_asc, Cooc12Pro_l1_des, Cooc12Pro_l1_max, Cooc12Pro_l2_asc, Cooc12Pro_l2_des, Cooc12Pro_l2_max, Cooc12Pro_l3_max, Cooc02ASM_mean, Cooc02ASM_skew, Cooc02ASM_max, Cooc02Con_mean, Cooc02Con_std, Cooc02Con_asc, Cooc02Con_des, Cooc02Con_max, Cooc02Con_l1_asc, Cooc02Con_l1_des, Cooc02Con_l2_asc, Cooc02Con_l2_des, Cooc02Con_l2_max, Cooc02Con_l3_asc, Cooc02Con_l3_des, Cooc02IDM_mean, Cooc02IDM_std, Cooc02IDM_asc, Cooc02IDM_des, Cooc02IDM_l1_asc, Cooc02IDM_l1_des, Cooc02IDM_l1_max, Cooc02IDM_l2_asc, Cooc02IDM_l2_des, Cooc02IDM_l2_max, Cooc02IDM_l3_asc, Cooc02IDM_l3_des, Cooc02IDM_l3_max, Cooc02Ent_skew, Cooc02Cor_mean, Cooc02Cor_std, Cooc02Cor_asc, Cooc02Cor_des, Cooc02Cor_max, Cooc02Cor_l1_asc, Cooc02Cor_l1_des, Cooc02Cor_l2_asc, Cooc02Cor_l2_des, Cooc02Cor_l3_asc, Cooc02Cor_l3_des, Cooc02Cor_l3_max, Cooc02Var_mean, Cooc02Var_skew, Cooc02Var_asc, Cooc02Var_des, Cooc02Var_max, Cooc02Var_l1_des, Cooc02Var_l1_max, Cooc02Var_l2_asc, Cooc02Var_l2_max, Cooc02Var_l3_max, Cooc02Sav_std, Cooc02Sav_skew, Cooc02Sav_asc, Cooc02Sav_max, Cooc02Sen_skew, Cooc02Sen_max, Cooc02Den_mean, Cooc02Den_asc, Cooc02Den_des, Cooc02Den_max, Cooc02Den_l1_asc, Cooc02Den_l1_des, Cooc02Den_l2_asc, Cooc02Den_l2_des, Cooc02Den_l3_des, Cooc02Dva_mean, Cooc02Dva_std, Cooc02Dva_asc, Cooc02Dva_des, Cooc02Dva_l1_asc, Cooc02Dva_l1_des, Cooc02Dva_l2_asc, Cooc02Dva_l2_des, Cooc02Dva_l3_asc, Cooc02Dva_l3_des, Cooc02Sva_std, Cooc02Sva_max, Cooc02Sva_l2_max, Cooc02f13_skew, Cooc02f13_des, Cooc02f13_max, Cooc02f13_l1_asc, Cooc02f13_l3_asc, Cooc02Sha_mean, Cooc02Sha_std, Cooc02Sha_asc, Cooc02Sha_des, Cooc02Sha_max, Cooc02Sha_l1_asc, Cooc02Sha_l1_des, Cooc02Sha_l1_max, Cooc02Sha_l2_asc, Cooc02Sha_l2_des, Cooc02Sha_l2_max, Cooc02Sha_l3_asc, Cooc02Sha_l3_des, Cooc02Sha_l3_max, Cooc02Pro_mean, Cooc02Pro_std, Cooc02Pro_skew, Cooc02Pro_asc, Cooc02Pro_des, Cooc02Pro_max, Cooc02Pro_l1_asc, Cooc02Pro_l1_des, Cooc02Pro_l1_max, Cooc02Pro_l2_asc, |

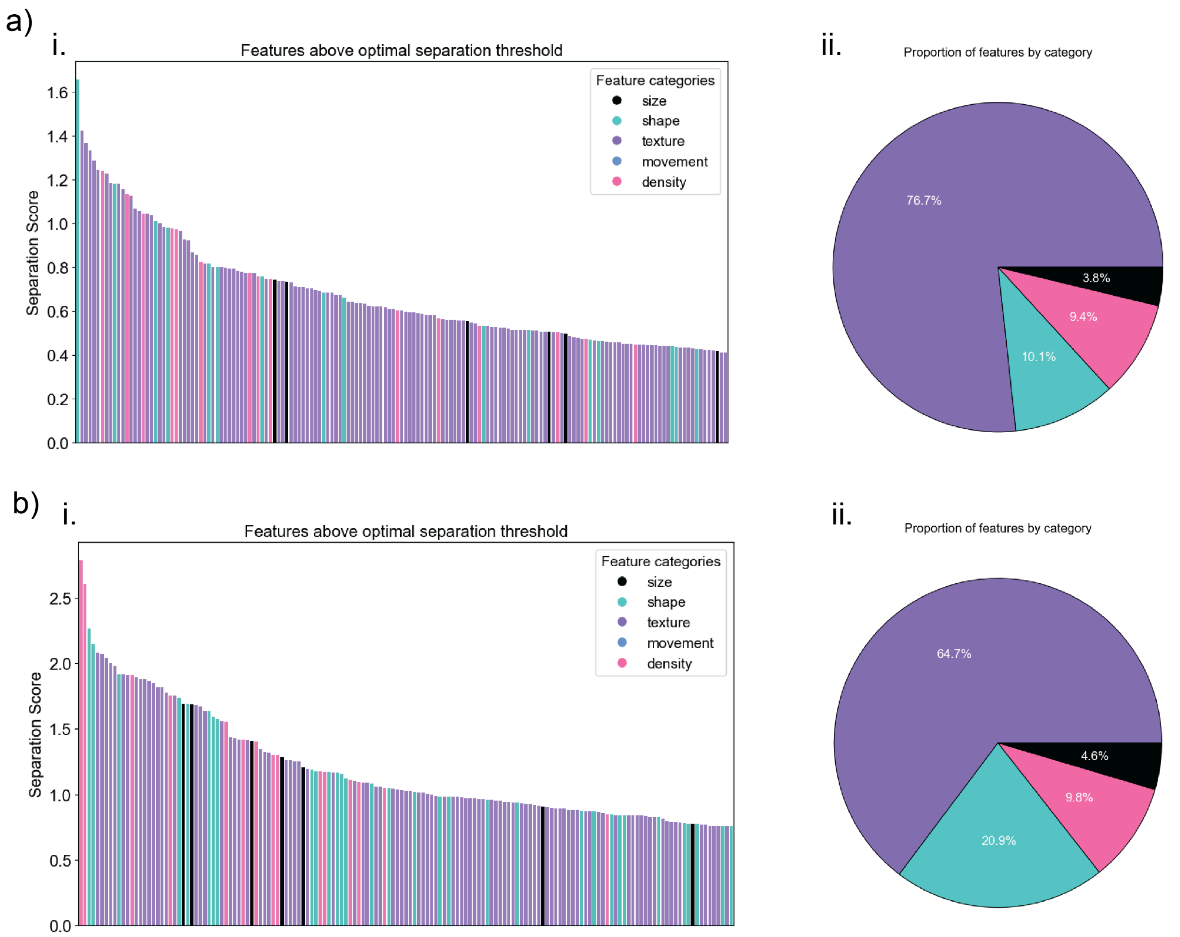


**Supplementary Figure 1. Discriminatory features distinguishing MDA-MB-231 vs. CAF and MCF-7 vs. CAF. a)** i. Bar plot of separation scores for MDA-MB-231 vs. CAF. ii. Pie chart showing the proportion of feature categories above the optimal separation threshold for MDA-MB-231 vs. CAF. **b)** i. Bar plot of separation scores for MCF-7 vs. CAF. ii. Pie chart showing the proportion of feature categories above the optimal separation threshold for MCF-7 vs. CAF.

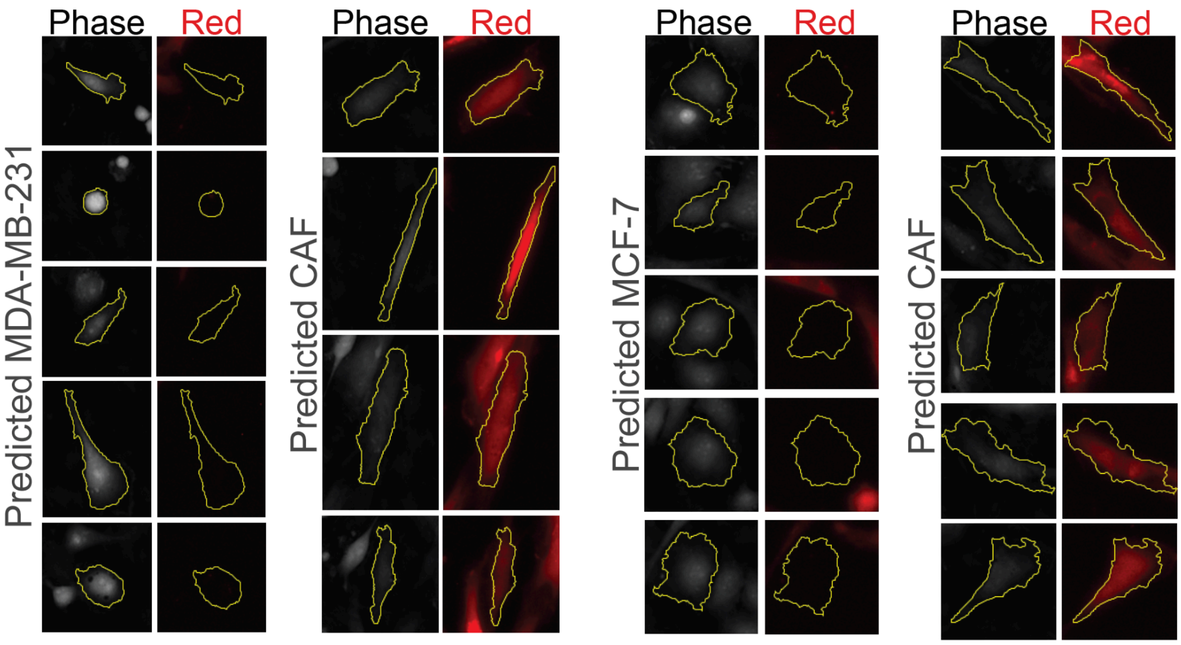


**Supplementary Figure 2. Representative microscopy images of cells classified as MDA-MB-231, MCF-7 and CAF.** Cells classified by the two models: the left panels display predictions from the MDA-MB-231 vs. CAF classifier (MDA-MB-231 or CAF), and the right panels display predictions from the MCF-7 vs. CAF classifier (MCF-7 or CAF).

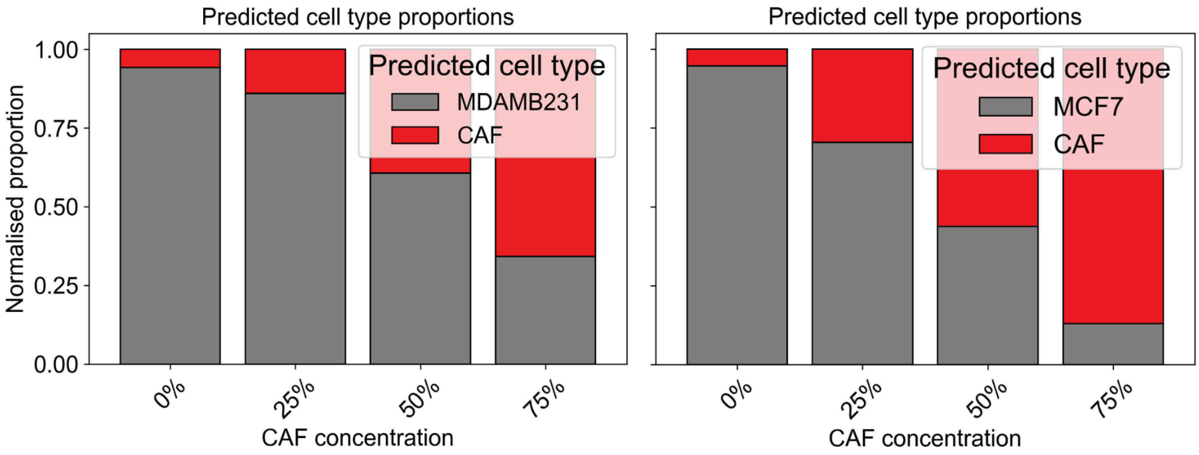


**Supplementary Figure 3. Predicted cell type proportions across CAF concentrations.** Predicted cell type labels for co-culture data sets based on k-means clustering and centroid matching. The proportion of predicted CAFs increased with seeding concentration: for MDA-MB-231 co-cultures, 6%, 14%, 40%, and 66% CAFs were identified at 0%, 25%, 50%, and 75% seeding, respectively; for MCF-7 co-cultures, the corresponding proportions were 5%, 30%, 56%, and 87%.

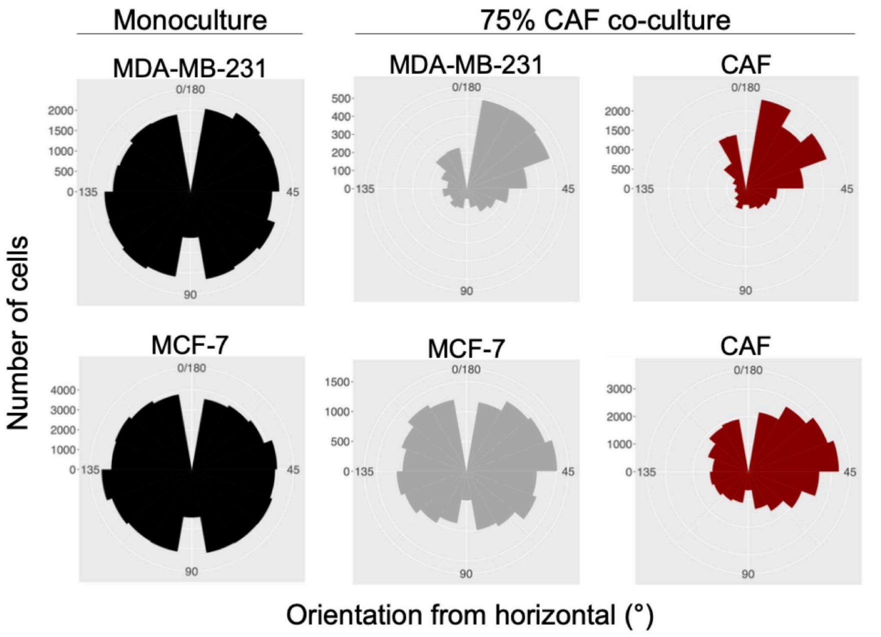


**Supplementary Figure 4. CAF co-culture induces alignment of MDA-MB-231 and MCF-7 cells with CAF orientation.** Rose diagrams to present orientation of cells from the main horizontal (classed as 0°). Diagrams demonstrate no bias towards any orientation within monoculture data sets but show increased alignment of co-cultured MDA-MB-231 and MCF-7 cells that correlate with the orientation of CAFs.

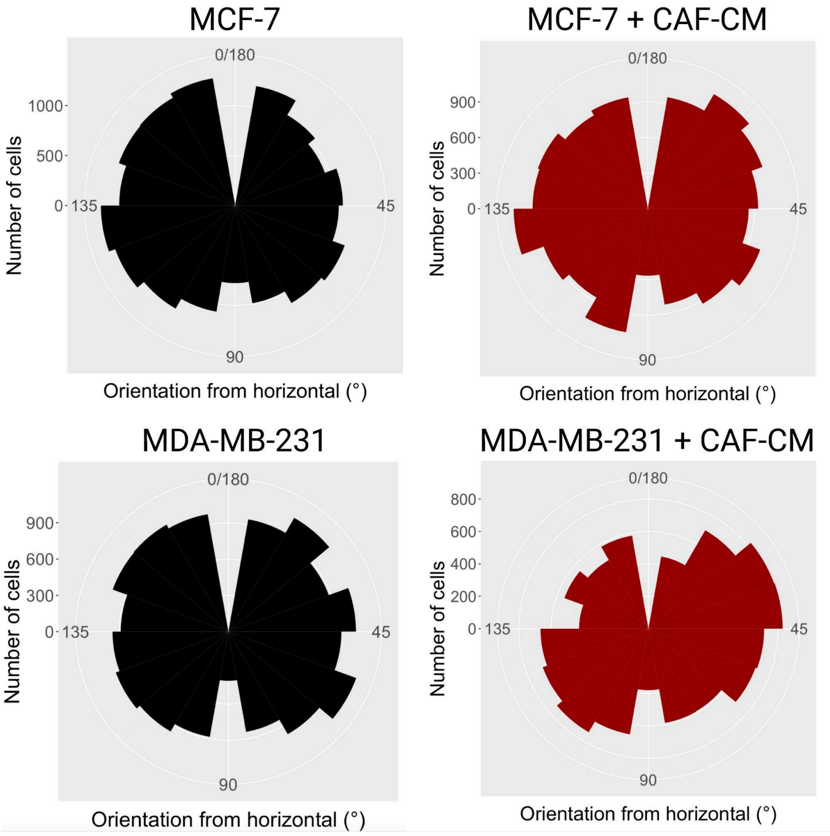


**Supplementary Figure 5. Culture in CAF-CM induces alignment of MDA-MB-231, but not MCF-7 cells.** Rose diagrams demonstrating no obvious alignment of MCF-7 cells following culture in CAF-CM, but collective alignment of a subset of MDA-MB-231 cells with bias towards certain orientations away from the horizontal.

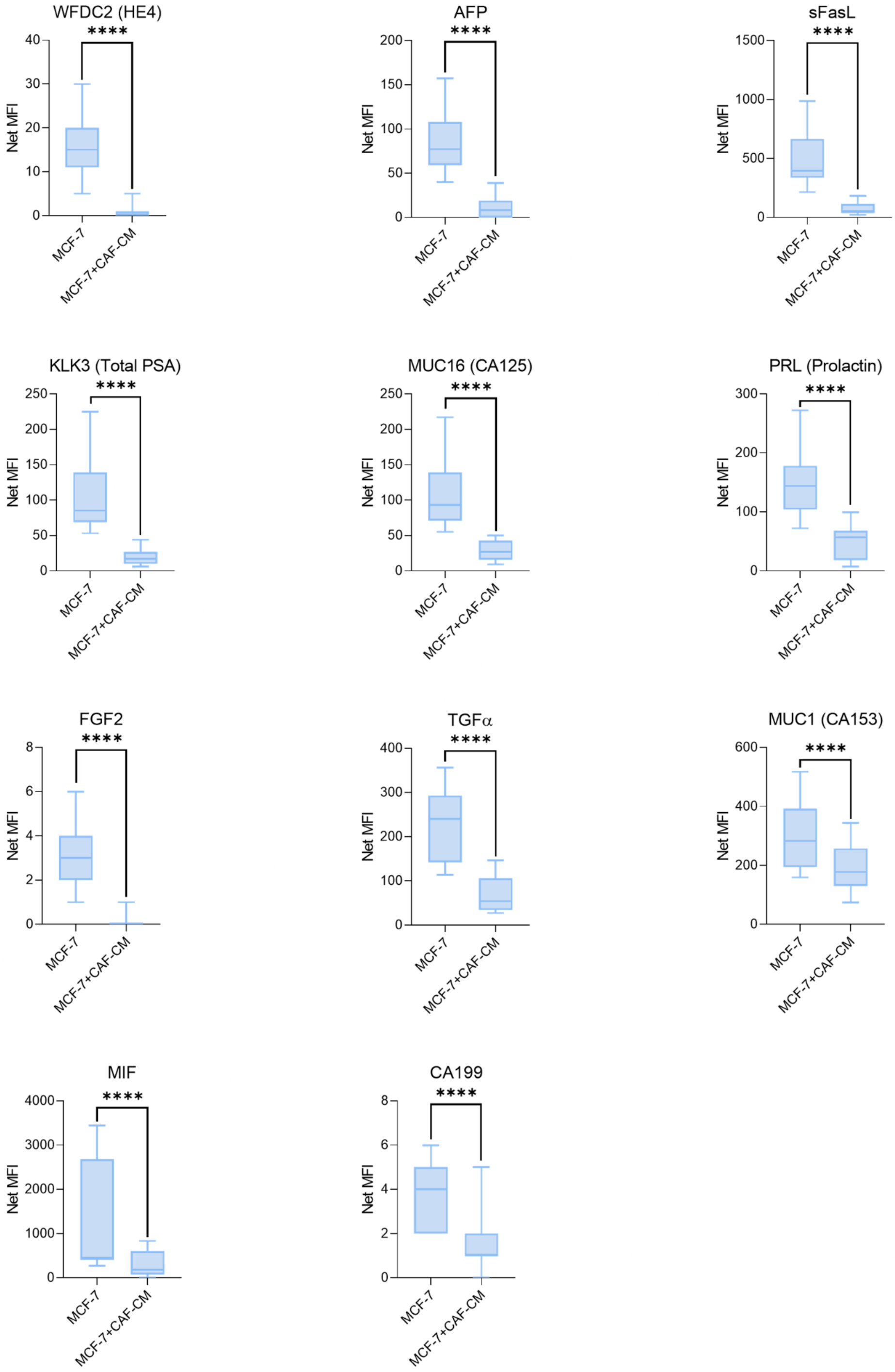


**Supplementary Figure 6. CAF-CM significantly reduces EMT-associated analytes in MCF-7 medium.** The addition of CAF-CM significantly decreased the presence of HE4, AFP, sFasL, Total PSA, CA125, Prolactin, FGF2, TGFa CA153, MIF and CA199in MCF-7 cells (N = 3).
